# The Functional Order (FunOrder) tool – Identification of essential biosynthetic genes through computational molecular co-evolution

**DOI:** 10.1101/2021.01.29.428829

**Authors:** Gabriel A. Vignolle, Denise Schaffer, Robert L. Mach, Astrid R. Mach-Aigner, Christian Derntl

## Abstract

Secondary metabolites (SMs) are a vast group of compounds with different structures and properties. Humankind uses SMs as drugs, food additives, dyes, and as monomers for novel plastics. In many cases, the biosynthesis of SMs is catalysed by enzymes whose corresponding genes are co-localized in the genome in biosynthetic gene clusters (BGCs). Notably, BGCs may contain so-called gap genes, that are not involved in the biosynthesis of the SM. Current genome mining tools can identify BGCs but they have problems with distinguishing essential genes from gap genes and defining the borders of a BGC. This can and must be done by expensive, laborious, and time-consuming comparative genomic approaches or co-expression analyses. In this study, we developed a novel tool that allows automated identification of essential genes in a BGC based solely on genomic data. The Functional Order (FunOrder) tool – Identification of essential biosynthetic genes through computational molecular co-evolution – searches for co-evolutionary linked genes in the BGCs. In light of the growing number of genomic data available, this will contribute to the studies of BGCs in native hosts and facilitate heterologous expression in other organisms with the aim of the discovery of novel SMs, including antibiotics and other pharmaceuticals.

## INTRODUCTION

Secondary metabolites (SMs) are a diverse group of compounds with a plethora of different chemical structures and properties. SMs are found in all domains of life, but are predominantly studied in bacteria, fungi, and plants (1). SMs are not necessary for the basic survival and growth of an organism but can be beneficial under certain conditions. For example, pigments help to sustain radiation, antibiotics help in competitive situations, and toxins can serve as defensive compounds or as virulence factors (2,3). Notably, many SMs are used by humankind as drugs and pharmaceuticals, pigments and dyes, sweeteners and flavours, and most recently also as precursors for the synthesis of plastics (4). The study of the secondary metabolism holds the promise for novel antibiotics, pharmaceuticals and other useful compounds (5).

A major hinderance in the discovery of yet undescribed SMs is the fact that most SMs are not produced under standard laboratory conditions, as they do not serve a purpose for the organisms then. Currently, different strategies are followed to circumvent this problem (6,7). Untargeted approaches aim to induce the expression of any SM. To this end, biotic and abiotic stresses are applied, or global regulators and regulatory mechanisms are manipulated (8). These strategies may lead to the discovery of novel compounds, whose corresponding genes have to be identified subsequently by time-consuming and expensive methods (7). An extreme example are the aflatoxins, major food contaminants with serious toxicological effects (9). It took over 40 years from the discovery of the aflatoxins as the causal agent of “turkey X” disease in the 1950s (10) until the corresponding genes were finally described in 1995 (11). Targeted SM discovery approaches aim to induce the production of specific SMs by either overexpression genes in the native host or by heterologous expression in another organism (12). The targeted approaches, also called reverse strategy or bottom-up strategy allows a direct connection of SMs to the corresponding genes and does not rely on the inducibility of SM production in the native host. Inherently, the bottom-up approach is depending on modern genomics and accurate gene prediction tools (13).

In bacteria and fungi, the genes responsible for the biosynthesis of a certain SM are often co-localized in the genome, forming so called biosynthetic gene clusters (BGCs) (14). The BGCs consists of one or more core genes and several tailoring enzymes. The core genes are responsible for assembling the basic chemical scaffold, which is further modified by the tailoring enzymes yielding the final SM (15). Depending on the class of the produced SM, the core genes differ. In bacteria and fungi, the main SM classes are polyketides (e.g. the cholesterol-lowering drug lovastatin (16) and the mycotoxin aflatoxin (9)) and non-ribosomal peptides (e.g. the immunosuppressant cyclosporine (17) and the antibiotic penicillin (18)), with polyketide synthases (PKS) or non-ribosomal peptide synthetases (NRPS) as core enzymes, respectively. Other SM classes are terpenoids, alkaloids, melanins (19,20), and ribosomally synthesized and posttranslationally modified peptides (RiPPs) (21,22), whose corresponding genes may also be organized in BGCs. Notably, BGCs contain not only genes necessary for the production of a SM, but also so-called gap genes that are not involved in the biosynthesis of the SM (Fig. 1).

**Figure 1.**
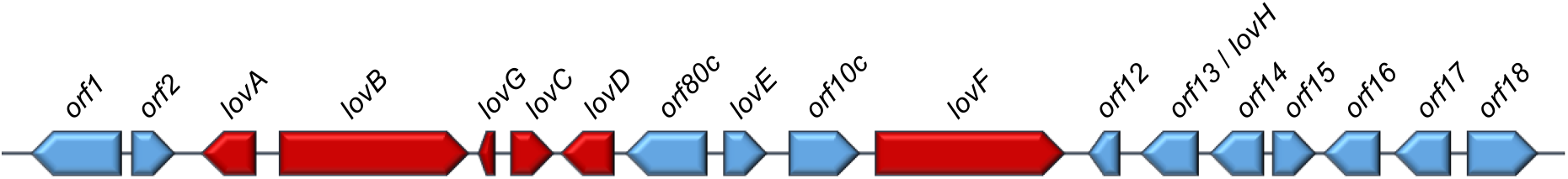
Schematic representation of the Lovastatin BGC from *Aspergillus terreus* (lov). In red the necessary genes for SM production and in blue the genes not involved in the biosynthetic pathway.

As mentioned, the bottom-up approach for SM discovery is depending on modern genomics and the accurate prediction of genes and BGCs. Each important gene missing in the prediction is detrimental for obvious reasons, whereas each unnecessarily considered gap gene and gene outside the BGC make the study of a BGC more complicated and complex, and the construction and transformation processes for heterologous expression more challenging. Currently, several BGC prediction tools are available for fungi and bacteria. For fungi, some prominent tools for genome mining are antiSMASH (23), Cassis (24) which has been integrated into antiSMASH, SMURF (25) and DeepBGC, a recently developed algorithm based on deep neural networks (26). For bacteria, the EvoMining tool predicts BGCs by searching for duplicates of primary metabolism enzymes (27). These tools are effective and successful in finding and predicting BCGs based solely on genomic data, but they do not predict gap genes. A further challenging task is the accurate determination of the borders of a BGC; the BGC prediction tools often contradict each other in this regard. These two problems can be solved by the analysis of transcriptome data, because the genes necessary for SM production within a BGC are co-expressed with each other but not with the gap genes and genes outside of the borders of the BGC (28). Notably, such methods demand the knowledge of expression conditions. Alternatively, the BGC can be analysed for co-evolution of the contained genes if no transcriptome data are available or the BGC is completely shut-off in the native host. A co-evolution analysis is a laborious and time-consuming task because a phylogenetic tree has to be calculated for each gene and then the trees compared to each other manually (29). However, it is an obvious advantage to have as much information as possible about a BGC before studying a BGC in the native host or performing heterologous expression for a bottom-up approach for SM discovery.

In this study we describe the Functional Order (FunOrder) tool – Identification of essential biosynthetic genes through computational molecular co-evolution, which identifies the relevant genes within a BGC based on the principle of co-evolution. We describe the workflow and demonstrate the applicability by an exemplary analysis of a well characterized BGC. We tested the robustness and the applicability of FunOrder by analysing empirically validated BGCs as positive controls, and randomly generated BGCs as negative controls and evaluated the FunOrder tool using stringent statistical tests. To our knowledge, FunOrder is the first program giving a prediction about core genes in fungal BGCs based solely on genomic data. Notably, the underlying strategy and workflow of FunOrder can be used to search for co-evolution in other phyla and for other objectives. FunOrder is available as command line version tool running on Linux and MacOS platforms.

## MATERIAL AND METHODS

### FunOrder - Workflow

The Functional Order (FunOrder) tool is written in the BASH (Bourn Again Shell) environment and includes all necessary subprograms. As BASH is the default shell-language of all Linux distributions and MacOS, FunOrder can run on these two operation systems. The only two dependencies are RAxML (Randomized Axelerated Maximum Likelihood) (30) and the EMBOSS (The European Molecular Biology Open Software Suite) package (31). The overall workflow is depicted in Figure 2. The FunOrder tool is deposited in the GitHub repository https://github.com/gvignolle/FunOrder. Notably, FunOrder includes scripts adapted to the use on servers and for the integration in various pipelines.

**Figure 2.**
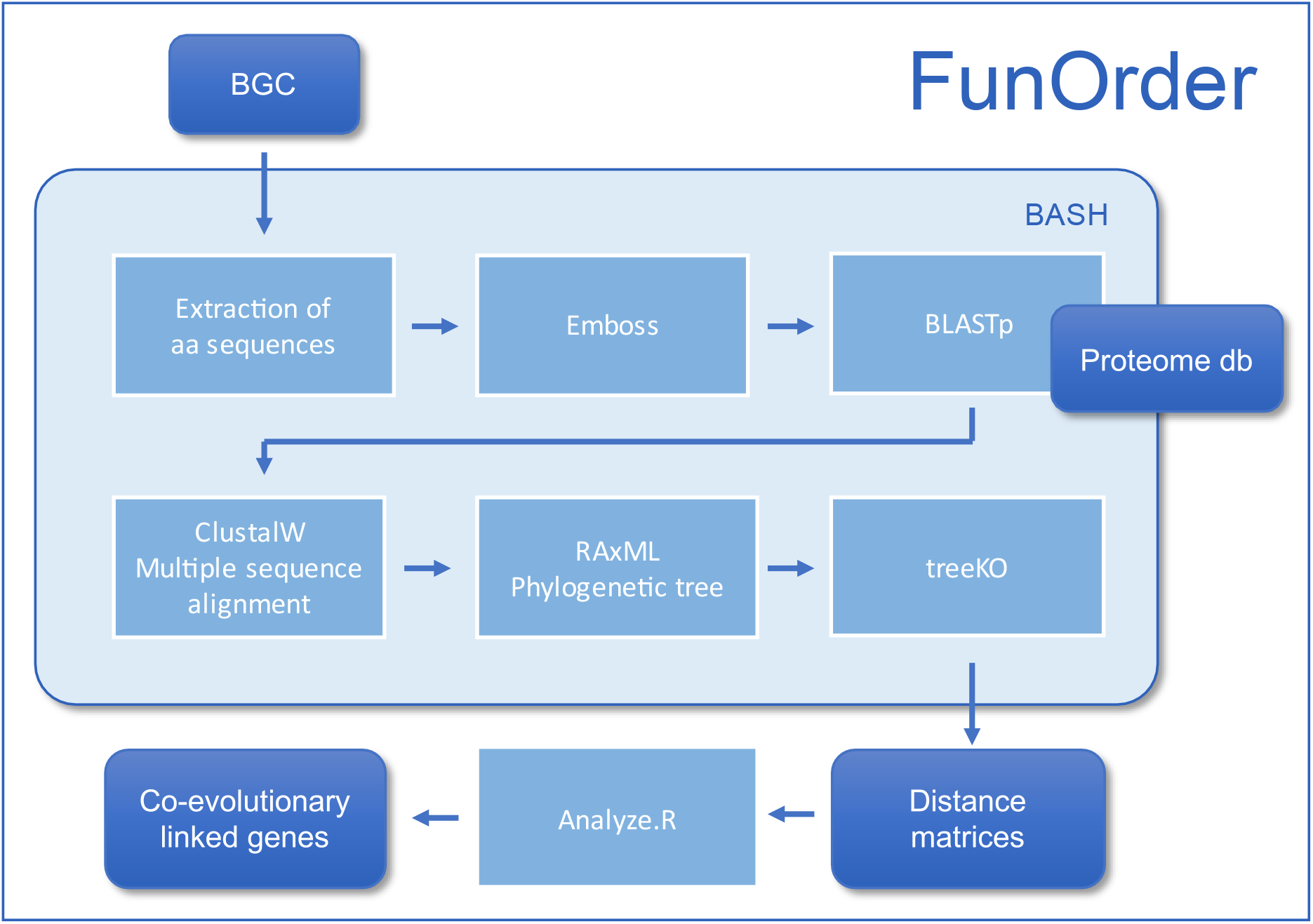
Schematic representation of the workflow of FunOrder.

The software input files are BGCs with gene translations in genbank file format or fasta format. In case a genbank file is provided, a python script (Genbank to FASTA by Cedar McKay and Gabrielle Rocap, University of Washington) is called to extract the amino acid sequence of the genes in the BGC and create a fasta file. The multi-fasta file is then split into individual fasta files each containing a single protein sequence. These are placed in a subfolder created for the analysis of the BGC. Each file is named either after the position of the gene in the BGC or after the respective protein sequence description. This varies from the input file and the varying annotations used. Each header of the query sequences is tagged with the identifier “query” at the beginning of the header. The individual sequences are compared to our manually curated database of fungal proteomes (Table 1) by a sequence similarity search using blastp 2.8.1+ (Protein-Protein BLAST) (32). The output of this search is saved in a file with the “.tab” extension. Additionally, an optional remote search of the non-redundant National Center for Biotechnology Information (NCBI) protein database can be performed, yielding a file with the “ncbi.tab” extension. This allows a manual preliminary analysis of the input sequences and supports subsequent annotations of the BGCs.

Next, the top 20 results of the database search of our manually curated fungal proteome database are extracted and combined with the query sequence for each gene. A custom Perl script removes potential duplicate entries. Using emma, a multiple sequence alignment of these protein sequences is calculated based on the ClustalW (33) algorithm, and a dendrogram computed. Based on the multiple sequence alignment 100 rapid Bootstraps and a subsequent search for the best-scoring maximum likelihood (ML) tree are performed using RAxML (Randomized Axelerated Maximum Likelihood) (30). The phylogenetic trees are computed using the LG amino acid substitution model. Furthermore, a standard ascertainment bias correction by Paul O. Lewis is performed. At this stage, we have obtained a phylogenetic tree (within the context of our manually curated database) for each gene of the input BGC.

To estimate if and to what extent the different genes within a BGC are co-evolved, the strict distance and speciation distance among the ML trees of the individual genes are calculated using the TreeKO algorithm (34). This tool was designed for automated tree comparison and was already suggested to be used for the detection of co-evolution in protein families (34). The tool compares the topology of different trees; a distance of 0 in both distance measures represents identical trees. In this context, a higher similarity between the different trees of the individual genes points towards a shared evolution. The strict distance is a weighted Robinson-Foulds (RF) distance measure that penalizes dissimilarities in evolutionarily important events such as gene losses and gene duplications. Furthermore the strict distance has been suggested to be more significant in the detection of co-evolution (34). In contrast, the speciation or evolutionary distance is computed without taking evolutionary exceptions, such as duplication events or different species content of the two compared trees, into account and infers shared “speciation history” based solely on topology without considering branch lengths and only considering shared species of the compared trees. Therefore, an evolutionary distance of 0 does not necessarily describe identical trees but shared “speciation history” of shared species. The comparison and analysis of the two distance measures is made possible through a custom BASH script and a special identifier introduced in the fungal proteome database. This enables a more comprehensive picture of the potential co-evolution within the analysed BGC. All pairwise strict and evolutionary distances are combined into matrices which are used as input for an R script (35–39).

First, the strict and evolutionary distances are summed up to a third combined distance measure. Then, the three distance matrices are visualized as heatmaps with a dendrogram computed with the complete linkage method, to find similar clusters. Next, the Euclidean distance within the matrices is computed and clustered using Ward’s minimum variance method aiming at finding compact spherical clusters, with the implemented squaring of the dissimilarities before cluster updating, for each of the three distance matrices separately, with scaled and unscaled input data (40). Lastly, a principal component analysis (PCA) is performed on each distance matrix and the score plot of the first two principal components visualized, respectively. These outputs enable the adoption of a larger view on the distance measures and thereby allow the analysis of co-evolution within the BGC from different perspectives.

### Manual curated fungal proteome database

The manual curated database used for sequence similarity search consists of 134 fungal proteomes, containing mainly ascomycetes and two basidiomycetes as potential outgroup (Table 1). The sequences were downloaded from the National Center for Biotechnology Information (NCBI) database and the Joint Genome Institute (JGI) (41). A short identifier, unique in the database for each proteome, was introduced to enable multiple pairwise tree comparisons by the treeKO application. A custom Perl script was used for removing duplicated entries in the database. The database is deposited in the GitHub repository https://github.com/gvignolle/FunOrder.

**Table 1.** Fungal proteomes included in the manually curated database. The sequences were downloaded from the National Center for Biotechnology Information (NCBI) database or the Joint Genome Institute (JGI). The identifiers were used in the FunOrder tool.

| Organism | Source Database | Identifier | Reference |
| --- | --- | --- | --- |
| *Acremonium chrysogenum* | JGI | AcCh | (42) |
| *Alternaria alternata* | NCBI | AlAl | (43) |
| *Alternaria arborescens* | NCBI | AlAr | (44) |
| *Alternaria gaisen* | NCBI | AlGa | (45) |
| *Alternaria sp.* MG1 | NCBI | AlSp | (46) |
| *Alternaria tenuissima* | NCBI | AlTe | (45) |
| *Amanita muscaria* | NCBI | AmMu | (47) |
| *Amorphotheca resinae* | JGI | AmRe | (48) |
| *Arthrotrys oligospora* | JGI | ArOl | (49) |
| *Arthroderma benhamiae* | JGI | ArBe | (50) |
| *Ascobolus immersus* | JGI | AsIm | (51) |
| *Aspergillus costaricensis* | NCBI | AsCo | (52) |
| *Aspergillus fijiensis* | NCBI | AsFi | (52) |
| *Aspergillus flavus* | NCBI | AsFl | (53) |
| *Aspergillus fumigatus* | NCBI | AsFu | (54) |
| *Aspergillus homomorphus* | NCBI | AsHo | (52) |
| *Aspergillus ibericus* | NCBI | AsIb | (52) |
| *Aspergillus japonicus* | NCBI | AsJa | (52) |
| *Aspergillus niger* | NCBI | AsNi | (55) |
| *Aspergillus oryzae* | NCBI | AsOr | (56) |
| *Aspergillus phoenicis* | NCBI | AsPh | (57) |
| *Aspergillus terreus* | NCBI | AsTe | (58) |
| *Blumeria graminis* | JGI | BlGr | (59) |
| *Botryosphaeria dothidea* | JGI | BoDo | (60) |
| *Botrytis cinerea* | NCBI | BoCi | (61) |
| *Botrytis elliptica* | NCBI | BoEl | (62) |
| *Botrytis galanthina* | NCBI | BoGa | (62) |
| *Botrytis hyacinthi* | NCBI | BoHy | (62) |
| *Botrytis paeoniae* | NCBI | BoPa | (62) |
| *Botrytis porri* | NCBI | BoPo | (62) |
| *Botrytis tulipae* | NCBI | BoTu | (62) |
| *Cadophora* sp. | JGI | CaSp | (63) |
| *Capronia semiimmersa* | JGI | CaSe | (64) |
| *Chaetomium globosum* | JGI | ChGl | (65) |
| *Choiromyces venosus* | JGI | ChVe | (51) |
| *Cladonia grayi* | JGI | ClGr | (66) |
| *Cladophialophora bantiana* | JGI | ClBa | (64) |
| *Cladophialophora carrionii* | JGI | ClCa | (64) |
| *Cladophialophora immunda* | JGI | ClIm | (64) |
| *Cochliobolus heterostrophus* | JGI | CoHe | (67) |
| *Cochliobolus victoriae* | JGI | CoVi | (68) |
| *Colletotrichum nymphaeae* | JGI | CoNy | (69) |
| *Colletotrichum orchidophilum* | JGI | CoOr | (70) |
| *Colletotrichum salicis* | JGI | CoSa | (69) |
| *Colletotrichum simmondsii* | JGI | CoSi | (69) |
| *Colletotrichum tofieldiae* | JGI | CoTo | (71) |
| *Coniosporium apollinis* | JGI | CoAp | (64) |
| *Coniosporium apollinis* CBS 100218 | JGI | Capo | (64) |
| *Corynespora cassicola* | JGI | CoCa | (72) |
| *Daldinia eschscholzii* | JGI | DaEs | (73) |
| *Diaporthe ampelina* | JGI | DiAm | (74) |
| *Diplodia seriata* | JGI | DiSe | (74) |
| *Erysiphe necator* | JGI | ErNe | (75) |
| *Eutypa lata* | NCBI | EuLa | (76) |
| *Exophiala aquamarina* | JGI | ExAq | (64) |
| *Exophiala dermatitidis* | JGI | ExDe | (64) |
| *Exophiala oligosperma* | JGI | ExOl | (64) |
| *Exophiala spinifera* | JGI | ExSp | (64) |
| *Exophiala xenobiotica* | JGI | ExXe | (64) |
| *Fonsecaea monophora* | JGI | FoMo | (77) |
| *Fusarium fujikuroi* | NCBI | FuFu | (78) |
| *Fusarium graminearum* | NCBI | FuGr | (79) |
| *Fusarium oxysporum* | NCBI | FuOx | (80) |
| *Fusarium proliferatum* | NCBI | FuPr | (81) |
| *Fusarium pseudograminearum* | NCBI | FuPs | (82) |
| *Fusarium verticillioides* | NCBI | FuVe | (79) |
| *Gaeumannomyces graminis* | JGI | GaGr | (83) |
| *Glonium stellatum* | JGI | GlSt | (84) |
| *Hypoxylon* sp. EC38 | JGI | HyEC | (73) |
| *Hypoxylon* sp.CO27 | JGI | Hysp | (73) |
| *Magnaporthe grisea* | JGI | MaGr | (85) |
| *Magnaportheopsis poae* | JGI | MaPo | (83) |
| *Meliniomyces bicolor* | JGI | MeBi | (48) |
| *Meliniomyces variabilis* | JGI | MeVa | (48) |
| *Metarhizium acridum* | NCBI | MeAc | (86) |
| *Metarhizium album* | NCBI | MeAl | (87) |
| *Metarhizium anisopliae* | NCBI | MeAn | (87) |
| *Metarhizium brunneum* | NCBI | MeBr | (87) |
| *Metarhizium guizhouense* | NCBI | MeGu | (87) |
| *Metarhizium majus* | NCBI | MeMa | (87) |
| *Metarhizium rileyi* | NCBI | MeRi | (88) |
| *Metarhizium robertsii* | NCBI | MeRo | (86) |
| *Monacrosporium haptotylum* | JGI | MoHa | (89) |
| *Morchella importuna* | JGI | Molm | (90) |
| [*Nectria* ] *haematococca* | NCBI | NeHa | (91) |
| *Nectria haematococca* | JGI | NeHa | (91) |
| *Neurospora crassa* | JGI | NeCr2 | (92) |
| *Neurospora crassa* FGSC | JGI | NeCr | (93) |
| *Neurospora tetrasperma* | JGI | NeTe | (94) |
| *Oidiodendron maius* | JGI | OiMa | (47) |
| *Ophiostoma piceae* | JGI | OpPi | (95) |
| *Paecilomyces variotii* | JGI | PaVa | (96) |
| *Panaeolus cyanescens* | NCBI | PaCy | (97) |
| *Paracoccidioides brasiliensis* | JGI | PaBr | (98) |
| *Penicillium camemberti* | NCBI | PeCa | (99) |
| *Penicillium chrysogenum* | NCBI | PeCh | (100) |
| *Penicillium digitatum* | NCBI | PeDi | (101) |
| *Penicillium expansum* | NCBI | PeEx | (102) |
| *Penicillium nalgiovense* | NCBI | PeNa | (103) |
| *Penicillium oxalicum* | NCBI | PeOx | (104) |
| *Penicillium roqueforti* | NCBI | PeRo | (99) |
| *Penicillium rubens Wisconsin* | NCBI | PeRu | (105) |
| *Penicillium vulpinum* | JGI | PeVu | (103) |
| *Periconia macrospinoso* | JGI | PeMa | (63) |
| *Pestalotiopsis fici* | NCBI | PeFi | (106) |
| *Phaeoacremonium aleophilum* | JGI | PhAl | (107) |
| *Phaeomoniella chlamydospora* | JGI | PhCh | (74) |
| *Phialocephala scopiformis* | JGI | PhSc | (108) |
| *Pneumocystis jirovecii* | JGI | PnJi | (109) |
| *Pseudogymnoascus destructans* | JGI | PsDe | (110) |
| *Pseudomassariella vexata* | JGI | PsVe | (111) |
| *Rhizoctonia solani* | NCBI | RhSo | (112) |
| *Saccharomyces arboricola* | NCBI | SaAr | (113) |
| *Saccharomyces cerevisiae* | NCBI | SaCe | (114) |
| *Terfezia boudieri* | JGI | TeBo | (51) |
| *Tolypocladium ophioglossoides* | NCBI | ToOp | (115) |
| *Tolypocladium paradoxum* | NCBI | ToPa | (116) |
| *Trichoderma arundinaceum* | NCBI | TrAr | (117) |
| *Trichoderma asperellum* | NCBI | TrAs | (118) |
| *Trichoderma atroviride* | NCBI | TrAt | (119) |
| *Trichoderma citrinoviride* | NCBI | TrCi | (118) |
| *Trichoderma harzianum* | NCBI | TrHa | (120) |
| *Trichoderma longibrachiatum* | NCBI | TrLo | (121) |
| *Trichoderma reesei* | NCBI | TrRe | (122) |
| *Trichoderma virens* | NCBI | TrVi | (119) |
| *Trichophyton rubrum* | JGI | TrRu | (123) |
| *Tuber aestivum* var. *urcinatum* | JGI | TuAe | (51) |
| *Tuber magnatum* | JGI | TuMa | (51) |
| *Venturia inaequalis* | JGI | VeIn | (124) |
| *Verruconis gallopava* | JGI | VeGa | (64) |
| *Verticillium dahliae* | JGI | VeDa | (125) |
| *Xylona heveae* | JGI | XyHe | (126) |
| *Zymoseptoria brevis* | JGI | ZyBr | (127) |
| *Zymoseptoria pseudotritici* | JGI | ZyPs | (128) |

### Compilation of benchmark biosynthetic gene clusters

To test and evaluate the applicability of FunOrder, we compiled negative and positive control BGCs (sequences deposited in the GitHub repository https://github.com/gvignolle/FunOrder). The first set of negative controls were 42 completely randomly generated synthetic BGCs, which were created with a custom BASH script. Therein, ATGC strings of random composition and length were translated to an amino acid string using transeq from the EMBOSS package and the asterisks were removed. The second set of negative controls were 60 random BGCs which were created by subsampling randomly the fungal proteome database with a Perl script from the MEME suit (129). For each random BGC a different seed number was given to guarantee non repetitive BGCs, each random BGC contained 3-10 randomly chosen protein sequences in a random order.

We used a set of 30 empirically well characterized BGCs from a broad range of different genera as positive controls (Table 2). The sequences were downloaded from NCBI or the MIBiG (Minimum information about a biosynthetic gene cluster) database (130). All BGCs were manually inspected for correctness and completeness based on the respective literature.

**Table 2.** Empirically characterized biosynthetic gene clusters used as positive controls.

| Product - BGC | Organism | MIBiG id | Reference(s) |
| --- | --- | --- | --- |
| 2-Pyridon-Desmethylobassianin (dmb) | *Beauveria bassiana* | BGC0001136 | (131) |
| Aflatoxin (afl) | *Aspergillus flavus* | BGC0000008 | (132,133) |
| Botrydial (bot) | *Botrytis cinera* | BGC0000631 | (134,135) |
| Cephalosporin (cef) | *Acremonium chrysogenum* | BGC0000317 | (136) |
| Compactin (mlc) | *Penicillium citrinum* | BGC0000039 | (137,138) |
| Cyclosporin (cyc2) | *Beauveria felina* | BGC0001565 | (17,139-141) |
| Destruxin (dtxs) | *Metarhizium robertsii* | BGC0000337 | (142) |
| Fumagillin (fma) | *Aspergillus fumigatus* | BGC0001067 | (143) |
| Fumitremorgin (ftm) | *Aspergillus fumigatus* | - | (144-147) |
| Fumonisin (fum1) | *Fusarium oxysporum* | BGC0000063 | (148) |
| Fumonisin (fum2) | *Fusarium verticilloides* | BGC0000062 | (149-156) |
| Fusaric acid (FUB) | *Fusarium fujikuroi* | - | (157) |
| Illicicolin H (ili) | *Neonectaria sp. DH2* | BGC0002035 | (158) |
| Leporin (lep) | *Aspergillus flavus* | BGC0001445 | (159) |
| Lovastatin (lov) | *Aspergillus terreus* | - | (16,58,160) |
| Mycophenolic acid (mpa1) | *Penicillium brevicompactum* | BGC0000104 | (161-166) |
| Mycophenolic acid (mpa2) | *Penicillium roqueforti* | BGC0001360 | (167) |
| Mycophenolic acid (mpa3) | *Penicillium roqueforti* | BGC0001677 | (168) |
| Paxillin (pax) | *Penicillium paxilli* | BGC0001082 | (169) |
| Penicillin (pen1) | *Penicillium chrysogenum* | BGC0000404 | (170) |
| Penicillin (pen2) | *Penicillium chrysogenum* | BGC0000405 | (18) |
| Pestheic acid (pta) | *Pestalotiopsis fici* | BGC0000121 | (171) |
| Pneumocandin (GL) | *Glaera lozoyensis* | BGC0001035 | (172-174) |
| Sorbicillinol (sor1) | *Penicillium rubens* | BGC0001404 | (175,176) |
| Sorbicillinol (sor2) | *Trichoderma reesei* | - | (177) |
| Tenellin (ten) | *Beauveria bassiana* | BGC0001049 | (178,179) |
| Terrein (ter) | *Aspergillus terreus* | BGC0000161 | (180) |
| Tetramic acid (tas) | *Hapsidospora irregularis* | - | (181) |
| Ustiloxin B (ust) | *Aspergillus flavus* | - | (182) |
| Xanthocillin (xan) | *Aspergillus fumigatus* | BGC0001990 | (183) |

### Performance evaluation

First, we performed a manual comparison of the phylogenetic trees for each control BGC and calculated a manual similarity measurement (manual evaluation measure - MEM) for each gene pair. Branch lengths, number of nodes, topology and shared leafs were considered (see Supplementary data Table S1 and S2). The measure ranges from 3 (same) to 0 (no shared leafs) (see Supplementary data “MEM_values.xlsx”). The MEM values were evaluate using thresholds of 1.2 and 2.0 for the negative control BGCs and for the positive control BGCs, respectively. These two thresholds were determined to reflect the biologically relevant evolution and to circumvent the selection bias introduced in the positive control BGCs (see Table S3). The MEM values were then compared to distance measures calculated by the treeKO algorithm (i.e., the strict distance, the evolutionary distance, and the combined distance measure) by exploratory data analysis. For the positive control BGCs, only those genes that had previously been described as necessary for the SM production were analysed. Based on these comparisons, we set threshold values for the treeKO analysis. Two genes are considered as co-evolved in a BGC, if the strict and evolutionary distance value is below 0.7 and the combined distance ≤ (0.6 * max value).

Next, we calculated three measures (two measures for the positive control BGCs and one for the negative control BGCs), to evaluate the performance of FunOrder regarding its capability to find coevolved genes (necessary for the biosynthesis of an SM), to distinguish them from not co-evolved (gap) genes, and to test FunOrder for its robustness, respectively. The previously set threshold values for the treeKO analysis was used to define which genes were detected as co-evolved by the FunOrder tool; genes had to have low distance values to each other (< 0.7 for strict and evolutionary distance and ≤ (0.6 * max value) for the combined distance) and/or cluster together in the PCAs, in the Ward’s minimum distance dendrograms and/or the heatmaps. Notably, the three measures are linked, meaning a single threshold for their subsequent evaluation can be used.

The measure to find the correct genes (find correct genes measure, FCGM) is a relative value that expresses how well the FunOrder tool identified genes that are necessary for the biosynthesis of a SM within the positive control BGCs. The FCGM was calculated using Equation 1, resulting in values between 0 and 1, with 1 representing a complete success in finding all necessary genes in the tested positive control BGC.

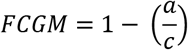

**Equation 1**. FCGM = find correct genes measure; a = number of genes necessary for the biosynthesis of a SM, that did not cluster with the other necessary genes in the FunOrder analysis; c = total number of genes necessary for the SM production in the BGC.

The error rate measure (ERM) is a relative value that expresses how well the FunOrder tool performed in finding the correct genes and differentiating them from the genes in a BGC not needed for the biosynthesis of a SM. The ERM was calculated using Equation 2, resulting in values between 0 and 1, with 1 representing no false classifications in the tested positive control BGC.

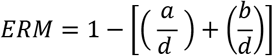

**Equation 2**. ERM = error rate measure; a = number of genes necessary for the biosynthesis of a SM, that did not cluster with the other necessary genes in the FunOrder analysis; b = number of gap genes, that clustered with the genes needed for the SM biosynthesis in the FunOrder analysis; d = number of analysed genes in the BGC.

The negative control value (NCV) is a relative value that expresses the robustness of the FunOrder tool. The NCV was calculated using Equation 3, resulting in values between 0 and 1, with 1 representing no detected co-evolved genes in the tested negative control BGC.

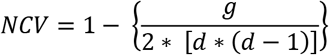

**Equation 3**. NCV = negative control value; d = number of genes in the BGC; g = number of strict distances < 0.7 and combined distances < (0.6 * max value) in all matrices.

Next, the obtained values for FCGM and ERM were classified as true positives (TP) or false negatives (FN), and the values for NCV were classified as true negative (TN) or false positives (FP). As the FCGM, the ERM, and the NCV are linked, a shared threshold was used for the classification. The threshold value was determined as the first quartile of all NCVs. This ensured an unbiased evaluation of the FunOrder tool. All values above the threshold were classified as true positives (TP) for the FCGM and the ERM and as true negatives (TN) for the NCV. All values below the threshold were classified as false negatives (FN) for the FCGM and the ERM and as false positives (FP) for the NCV. Finally, the results of the FunOrder analysis of the positive control BGCs (TP vs. FN) were assayed in two confusion matrices.

## RESULTS AND DISCUSSION

### Exemplary analysis of the Lovastatin BGC (lov)

FunOrder compares the genes in a BGC regarding their evolutionary background and calculates two distance matrices, the strict distance, and the evolutionary distance. These distance matrices are then combined, analysed, and visualized in different ways (Fig. 3, Supplementary data Fig. S1 – S7). To evaluate if and which genes are necessary for the biosynthesis of a SM, we searched for genes that were clustering together with the backbone producing genes (e.g., PKS or NRPS) in the visualisations. Genes clustering together can be considered as co-evolutionary linked genes, it is thus likely that the respective enzymes are part of the same biosynthetic pathway. We exemplary describe the analysis for the positive control Lovastatin BGC of *Aspergillus terreus* (lov, Fig. 3). This analysis strategy was used for all control BGC in this study and can be used for the analysis of yet uncharacterized BGCs.

**Figure 3.**
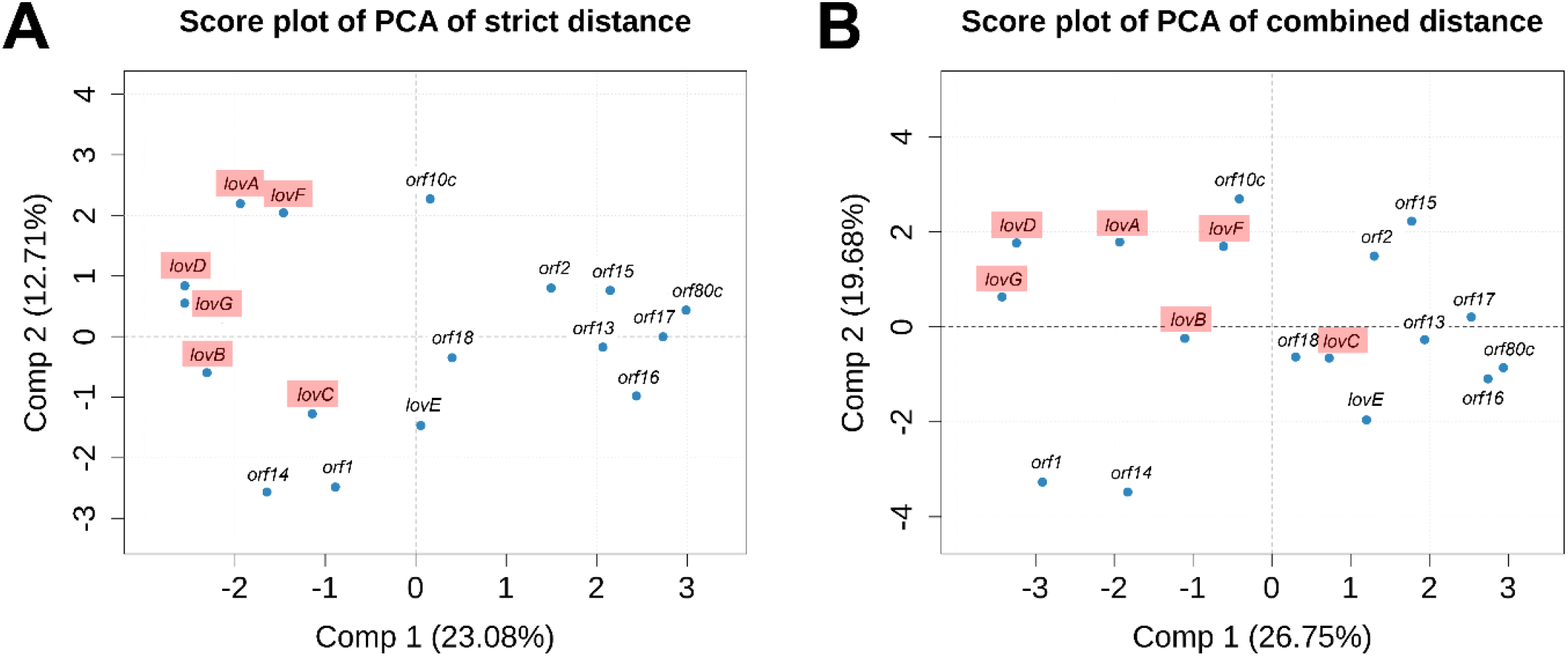
A A selection of the standard output of the FunOrder analysis of the Lovastatin BGC (lov, Fig. 1A). (**A**) Score plot of the first two principal components from a PCA performed on the combined distance matrix. (**B**) Score plot of the first two principal components from a PCA performed on the strict distance matrix. Gene necessary for the biosynthesis of lovastatin are highlighted in red.

For the evaluation and interpretation of the FunOrder results for the Lovastatin BGC (Fig. 3, Supplementary data Fig. S1 – S7), we first had a look at the numerical distance matrices (Supplementary data, Tables S5 – S7). Not all of the calculated values for the genes necessary for the biosynthesis of lovastatin (*lovA-D, F,G;* Fig. 1, red arrows) fell within the previously defined limit of < 0.7 for strict and evolutionary distance and ≤ (0.6 * max value) for the combined distance. In our experience, evaluating only the numerical values is not enough for a thorough analysis of a BGC and it is necessary to consider all provided visualisations for a thorough data interpretation (Fig. 3, Supplementary data Fig. S1 – S7). All genes necessary to produce lovastatin *(lovA-D, F, G)* formed distinct clusters in the heatmap, in the dendrograms and in the PCA of the strict distance (Supplementary data Fig. S1, S4; Fig. 3A). In the visualizations of the evolutionary distance, on the other hand, the necessary genes did not cluster together and were indistinguishable from the gap genes (Supplementary data Fig. S2, S5, S7). This results are expected, as the strict distance has been suggested to be more significant in the detection of co-evolution between protein families (34). Consequently, we recommend using the strict distance values as the basis for data interpretation. In our experience, it was often helpful to additionally take the combined distance values into consideration to get a more comprehensive picture of the BGC. As mentioned before, the combined distance also considers speciation history. In the case of the Lovastatin BGC, *orf10c* clustered together with *lovA, B, D, F, G* in the PCA of the combined distance, whereas *lovC* was not clustering with the other necessary genes when considering the speciation history (Fig. 3B). The gene *orf10c* encodes for an MFS (major facilitator superfamily) transporter that is necessary for export of lovastatin (16), it is thus not directly necessary for the biosynthesis of lovastatin, but important for a lovastatin-producing organism. Analogously, *lovE* did not cluster with the other genes in the PCA of the combined distance matrix, nor in any other distance measure used for the detection of co-evolution (Fig. 3 and Supplementary data Fig. S1-S7). Notably, LovE is a transcription factor and the main regulator of the lovastatin cluster (16) and thus not directly part of the biosynthetic pathway of lovastatin. The obtained results show how varying speciation history of different genes influences the FunOrder output and that both distances should be considered for evaluation.

The Lovastatin BGC is a clear example of the ability of FunOrder to discover the correct genes for SM production and to differentiate them from gap genes. In this case differentiating *orf1, orf2, orf10c, orf13, orf14, orf15, orf16, orf17, orf18, lovE* and *orf80c* from the needed *lovA, lovB, lovC, lovD, lovF* and *lovG*. This exemplary analysis further points out how we needed to adopt a wider view of the data to make a more precise decision on which genes are co-evolutionary linked. When considering only one output, one might get a distorted view of the analysed BGC. Notably, the more visualisations point towards a coevolution of two or more genes, the more reliable the prediction of FunOrder.

### Speed and scalability

As the manually curated proteome database contained only 134 fungal proteomes, we were able to use the blastp algorithm for sequence similarity search. The analysis of the Lovastatin BGC of *Aspergillus terreus* (lov) with 17 genes, took 1 h 19 m 48 sec real time using 22 threads on an Ubuntu Linux system with 128 GB DDR4 RAM, demonstrating that the analysis of such a large cluster as the Lovastatin cluster is fast and feasible. The number of threads can be defined, to increase the scalability and the overall performance of FunOrder.

### Performance evaluation

#### Negative control BGCs

FunOrder did not find any sequence similarities between the 42 synthetic BGCs and the manually curated fungal proteome database. This demonstrates the robustness of FunOrder towards non-biological random amino acid sequences. Consequently, the synthetic BGCs were not considered when calculating the confusion matrix to evaluate the performance of FunOrder.

The random BGCs, on the other hand, consist of randomly sampled genes from the fungal proteome database itself and represent a suitable negative control set where each gene has a biological background. For each of the random BGC the NCV was calculated as described in equation 3. A low NCV value indicates, that FunOrder detected co-evolutionary linked genes in the random BGC, a high value indicates that FunOrder could not detect linked genes. The NCV minimum value was 0.5 and the maximum 1, the first quartile was 0.66, the third quartile 0.83 and the median 0.75. Based on a Shapiro-Wilk normality test (alpha significance value 0.05) the NCVs can be considered normally distributed (Fig. 4A), demonstrating that the used set of BGCs was indeed random and suitable for the usage as negative controls. Based on the first quartile of the NCVs an unbiased threshold was set to 0.66 for the classification into true negatives (TN) or false positives (FP). 49 of the 60 negative control BGCs were above the set threshold and were considered as TN. 11 of the negative control BGCs were below the threshold and considered FP (Supplementary data, Table S4).

**Figure 4.**
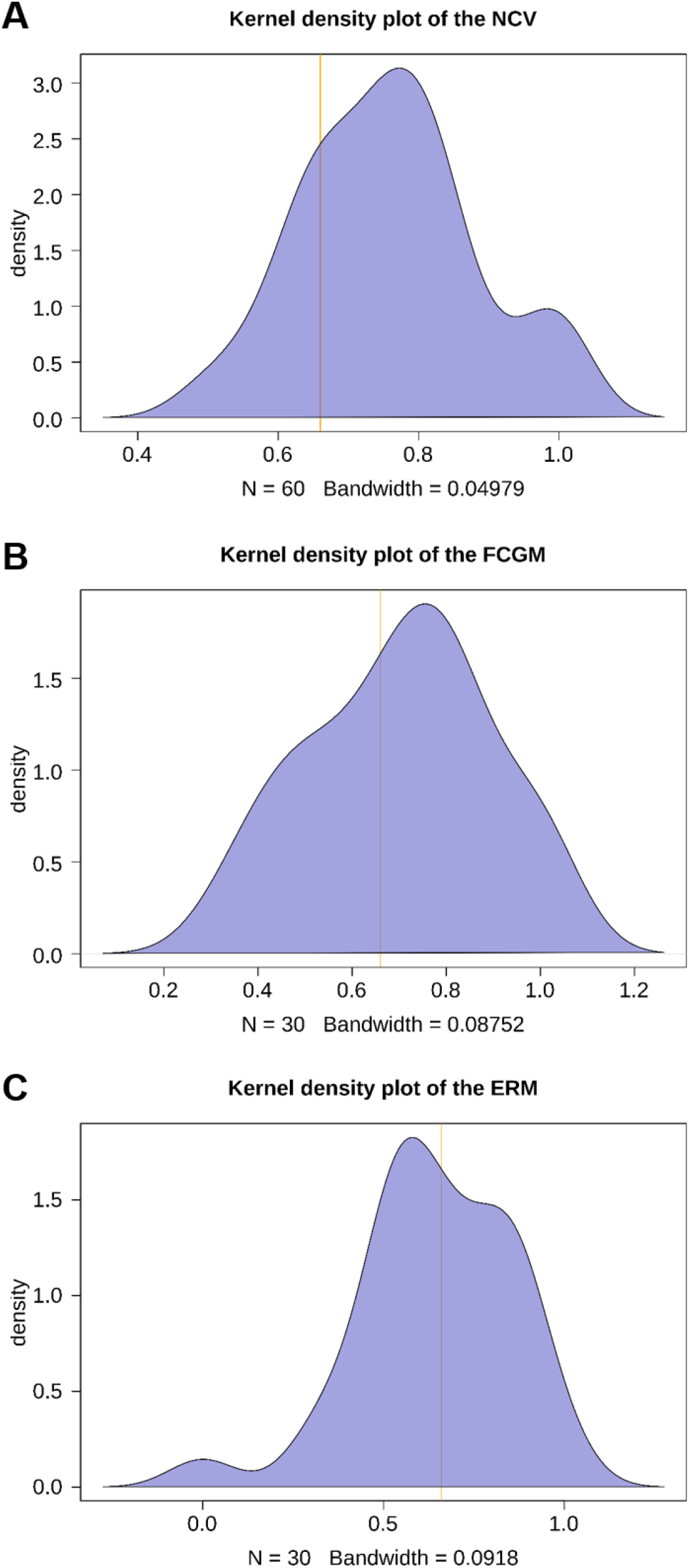
Kernel density plots of the (**A**) Negative control values (NCV) calculated based on equation 3, (**B**) the Find the correct genes measure (FCGM) calculated based on equation 1, and (**C**) the error rate measure (ERM) calculated based on equation 2. The vertical orange line indicates the 1^st^ quartile of the NCV and the chosen threshold.

#### Positive control BGCs

We used 30 empirically verified BGCs (Table 2) as positive controls for benchmarking the FunOrder tool. These BGCs were all previously characterized by *in vivo* molecular biology techniques, such as gene deletions or heterologous expression of the genes in different hosts. To evaluate how well the FunOrder tool performs in finding co-evolutionary linked genes and distinguish them from gap genes in positive control BGCs, we compared the results of the FunOrder analysis to the previously published experimental data. To this end, we first analysed each BGC with the FunOrder tool and looked for genes that are co-evolutionary linked with the core biosynthetic gene. These gene sets were considered as “detected” and compared to the empirically proven set of necessary genes. Based on these numbers, we calculated the FCGM and ERM (Equations 1 and 2) for each positive control BGC. A high FCGM value indicates that FunOrder was able identify a large proportion of the necessary genes in the analysed BGC. The FCGM minimum value was 0.33 and the maximum 1, the first quartile was 0.55, the third quartile 0.82 and the median 0.73. Based on a Shapiro-Wilk normality test (alpha significance value 0.05) the FCGMs can be considered normally distributed (Fig. 4B). 20 of the 30 positive control BGCs were above the set threshold of 0.66, these were considered TP. 10 of the positive control BGCs were below the threshold and were marked as FN (Table 3). A high ERM indicated a low error rate (undetected necessary genes or detected gap genes). The ERM minimum value was 0 and the maximum 1, the first quartile was 0.53, the third quartile 0.8 and the median 0.64. Based on a Shapiro-Wilk normality test (alpha significance value 0.05) the ERMs can be considered normally distributed (Fig. 4C). 15 of the 30 positive control BGCs were above the set threshold 0.66 and were considered as TP, whereas 15 of the positive control BGCs were below the threshold and considered FN (Table 3). As mentioned above a single threshold was chosen as the 1^st^ quartile of the NCV, to avoid a biased choice and a transparent evaluation. The previously exemplarily characterised Lovastatin BGC of *Aspergillus terreus* (lov) had a FCGM of 0.83 and an ERM of 0.88.

The numbers of TP and FN (Table 3) were used to create two confusion matrices (Fig. 5A and 5B), which were the basis for a thorough statistical testing of the applicability of the FunOrder tool (Table 4). We calculated the normalized Matthews correlation coefficient (normMCC) and other classical metrics and global metrics as indicated by Chicco and Jurman (184). To determine the degree of balance between positive and negative controls we calculated the *no-information error rate* ni, which is best for balanced test sets with the value 0.5. The evaluation showed that the ni value was at 0.67. This still allowed for the usage and confirmed the validity of the classical metrics such as F1 score and Accuracy. Based on the FCGM and the stringent classification threshold (0.66), FunOrder displays overall high metrics in identifying necessary genes in a BGC. Based on the ERM and the stringent classification threshold (0.66) the evaluation metrics can still be considered high regarding the ability of FunOrder to identify necessary genes and distinguish them from gap genes (Table 4).

**Figure 5.**
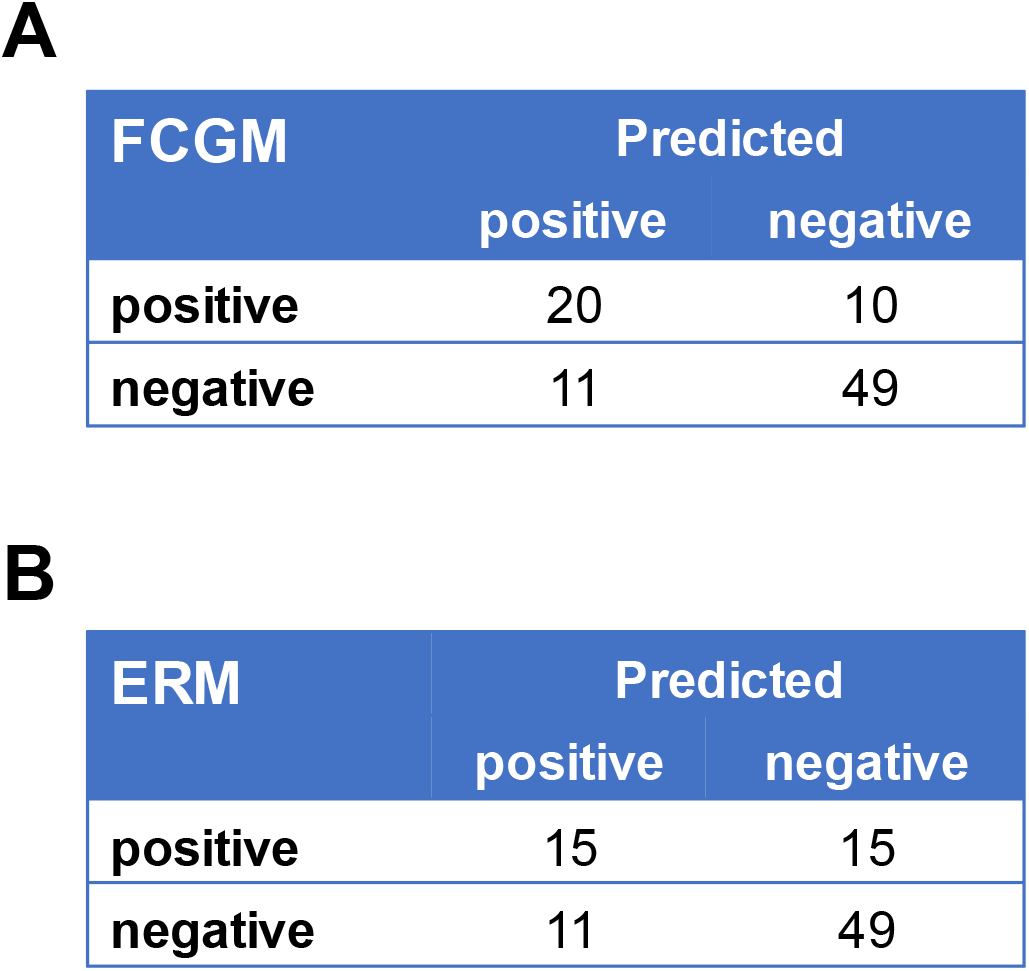
Confusion matrices calculated with (**A**) the Negative control values (NCV) and find the correct genes measure (FCGM) as basis for true negatives (TN), false positives (FP), true positives (TP) and false negatives (FN), (**B**) the NCV and error rate measure (ERM) as basis for TN, FP, TP, and FN.

**Table 3.** Analysis of the positive control BGCs and benchmarking of the FunOrder tool. The experimentally characterized BGCs were analysed with the FunOrder tool and the find correct genes measure (FCGM) and the error rate measure (ERM) were determined by manual comparison to the literature. Based on the thresholds of 0.66, the ERM and FCGM were classified as false negative (FN) or true positive (TP).

| Positive control BGC | Species | Genes | ERM | FCGM | (FN / TP)<br>ERM | (FN / TP)<br>FCGM |
| --- | --- | --- | --- | --- | --- | --- |
| 2-Pyridon-Desmethylbassianin (dmb) | *B. bassiana* | 4 | 0.50 | 0.50 | FN | FN |
| Aflatoxin (afl) | *A. flavus* | 23 | 0.35 | 0.43 | FN | FN |
| Botrydial (bot) | *B. cinera* | 7 | 0.86 | 0.67 | TP | TP |
| Cephalosporin (cef) | *A. chrysogenum* | 7 | 0 | 0.33 | FN | FN |
| Compactin (mlc) | *P. citrinum* | 9 | 0.55 | 0.67 | FN | TP |
| Cyclosporin (cyc2) | *B. felina* | 13 | 0.31 | 0.38 | FN | FN |
| Destruxin (dtxs) | *M. robertsii* | 21 | 0.62 | 1 | FN | TP |
| Fumagillin (fma) | *A. fumigatus* | 15 | 0.87 | 0.75 | TP | TP |
| Fumitremorgin (ftm) | *A. fumigatus* | 9 | 0.55 | 0.62 | FN | FN |
| Fumonisin (fum1) | *F. oxysporum* | 16 | 0.59 | 0.82 | FN | TP |
| Fumonisin (fum2) | *F. verticilloides* | 23 | 0.52 | 0.82 | FN | TP |
| fusaric acid (FUB) | *F. fujikuroi* | 15 | 0.87 | 0.92 | TP | TP |
| Illicolin H (ili) | *N. sp. DH2* | 6 | 0.67 | 0.75 | TP | TP |
| Leporin (lep) | *A. flavus* | 10 | 0.80 | 0.67 | TP | TP |
| Lovastatin (lov) | *A. terreus* | 17 | 0.88 | 0.83 | TP | TP |
| Mycophenolic acid (mpa1) | *P. brevicompactum* | 8 | 0.50 | 0.50 | FN | FN |
| Mycophenolic acid (mpa2) | *P. roqueforti* | 7 | 0.71 | 0.71 | TP | TP |
| Mycophenolic acid (mpa3) | *P. roqueforti* | 7 | 0.43 | 0.43 | FN | FN |
| Paxillin (pax) | *P. paxilli* | 8 | 1 | 1 | TP | TP |
| Penicillin (pen1) | *P. chrysogenum* | 15 | 0.87 | 1 | TP | TP |
| Penicillin (pen2) | *P. chrysogenum* | 15 | 0.93 | 1 | TP | TP |
| Pestheic acid (pta) | *P. fici* | 18 | 0.55 | 0.78 | FN | TP |
| Pneumocandin (GL) | *G. lozoyensis* | 25 | 0.68 | 0.80 | TP | TP |
| Sorbicillinol (sor1) | *P. rubens* | 7 | 0.57 | 0.57 | FN | FN |
| Sorbicillinol (sor2) | *T. reesei* | 15 | 0.80 | 0.80 | TP | TP |
| Tenellin (ten) | *B. bassiana* | 5 | 0.80 | 0.75 | TP | TP |
| Terrein (ter) | *A. terreus* | 10 | 0.57 | 0.86 | FN | TP |
| Tetramic acid (tas) | *H. irregularis* | 8 | 0.50 | 0.50 | FN | FN |
| Ustiloxin B (ust) | *A. flavus* | 19 | 0.68 | 0.54 | TP | FN |
| Xanthocillin (xan) | *A. fumigatus* | 7 | 0.71 | 0.67 | TP | TP |

**Table 4.** Metrics based on the two confusion matrices for FCGM (find correct genes measure) and ERM (error rate measure) (Fig.5). True positive rate (TPR), true negative rate (TNR), positive predictive value (PPV), negative predictive value (NPV), false negative rate (FNR), false positive rate (FPR), false discovery rate (FDR), false omission rate (FOR), Accuracy, F1 score, Matthews correlation coefficient (MCC) and normalized Matthews correlation coefficient (normMCC).

| **Metric** | **FCGM** | **ERM** |
| --- | --- | --- |
| TPR / Sensitivity | 0.667 | 0.5 |
| TNR / Specificity | 0.817 | 0.817 |
| PPV / Precision | 0.645 | 0.577 |
| NPV | 0.830 | 0.766 |
| FNR / miss rate | 0.333 | 0.5 |
| FPR / fall out | 0.183 | 0.183 |
| FDR | 0.355 | 0.423 |
| FOR | 0.170 | 0.234 |
| Accuracy | 0.767 | 0.711 |
| F1 Score | 0.656 | 0.536 |
| MCC | 0.4795 | 0.3294 |
| normMCC | 0.7397 | 0.6647 |

### Concluding remarks

In this study, we demonstrated that FunOrder can reliably find co-evolutionary linked genes in BGCs from ascomycetes. The basis but also limitation for FunOrder is the manually curated database. Here we used a set of ascomycete proteomes and were thus able to detect co-evolved genes in ascomycetes. The underlying strategy and workflow of FunOrder can be adapted to analysing genomic regions in other phyla, orders, or even kingdoms by using different databases. In case a larger database is integrated into FunOrder, alternative search algorithms, such as DIAMOND (185) or HMMER (186) (similarity search using hidden Markov models) might be used instead of blastp to enhance the performance.

Here, we further demonstrated that FunOrder can be applied to identify the genes of a BGC that are necessary for the biosynthesis of a SM. Based on a stringent threshold (Fig. 4), we calculated a normalized MCC of 0.7397 (Table 4) for the identification of the necessary genes. Further, with a normMCC of 0.6647 (Table 4) we were able to differentiate between necessary and gap genes in the analysed BGCs, this is also reflected in a high Accuracy of 71.1% (Table 4). The FunOrder tool is delivering data on co-evolution, that needs to be critically evaluated and interpreted keeping the biological background in mind. We introduced the previously discussed measures FCGM and ERM to enable a stringent and robust performance determination to prevent an overestimation of the evaluation metrics. In this study, we looked on genes that share the same or a similar evolutionary background with the core gene(s) of a BGC (e.g., PKS or NRPS-encoding genes). The optional remote search of the non-redundant NCBI protein database delivers further information on the analysed BGC, which can support the data interpretation. FunOrder is a fast and powerful tool that can support scientists to decide which genes of a BGC are promising study objects. The application of FunOrder is not limited to fungal BGC. It can be used for any applications, where information of a shared co-evolution can contribute to a better understanding, such as a genome wide analysis of co-evolving transcription factors or detection of functionally connected protein-protein interactions (29). As a future perspective, FunOrder might be even used for the analysis of total proteomes to detect evolutionary linked genes.

## DATA AVAILABILITY

The FunOrder tool, the relevant database, and the sequences and the FunOrder output of the negative and positive control BGCs are available in the GitHub repository (https://github.com/gvignolle/FunOrder).

## SUPPLEMENTARY DATA

Supplementary data file 1: MEM_values.xlxs

Supplementary data file 2: supplementary_tables_figures.pdf

## REFERENCES

1. Thirumurugan, D., Cholarajan, A., Raja, S.S.S. and Vijayakumar, R. (2018) In Vijayakumar, R. and Raja, S. S. S. (eds.), Secondary Metabolites - Sources and Applications. IntechOpen Limited, London, UK.

2. Malik, V.S. (1980) Microbial secondary metabolism. Trends in Biochemical Sciences, 5, 68–72.

3. Keller, N.P., Turner, G. and Bennett, J.W. (2005) Fungal secondary metabolism — from biochemistry to genomics. Nature Reviews Microbiology, 3, 937–947.

4. Alberti, F., Foster, G.D. and Bailey, A.M. (2017) Natural products from filamentous fungi and production by heterologous expression. Applied microbiology and biotechnology, 101, 493–500.

5. Newman, D.J., Cragg, G.M. and Kingston, D.G.I. (2015) In Wermuth, C. G., Aldous, D., Raboisson, P. and Rognan, D. (eds.), The Practice of Medicinal Chemistry (Fourth Edition). Academic Press, San Diego, pp. 101–139.

6. Brakhage, A.A. and Schroeckh, V. (2011) Fungal secondary metabolites - strategies to activate silent gene clusters. Fungal genetics and biology: FG & B, 48, 15–22.

7. Atanasov, A.G., Zotchev, S.B., Dirsch, V.M., Orhan, I.E., Banach, M., Rollinger, J.M., Barreca, D., Weckwerth, W., Bauer, R., Bayer, E.A. et al. (2021) Natural products in drug discovery: advances and opportunities. Nature Reviews Drug Discovery.

8. Wiemann, P. and Keller, N.P. (2014) Strategies for mining fungal natural products. Journal of industrial microbiology & biotechnology, 41, 301–313.

9. Kensler, T.W., Roebuck, B.D., Wogan, G.N. and Groopman, J.D. (2011) Aflatoxin: a 50-year odyssey of mechanistic and translational toxicology. Toxicol Sci, 120 Suppl 1, S28–48.

10. Blount, W. (1961) Turkey “X” disease. Turkeys, 9, 52–55.

11. Yu, J., Chang, P.K., Cary, J.W., Wright, M., Bhatnagar, D., Cleveland, T.E., Payne, G.A. and Linz, J.E. (1995) Comparative mapping of aflatoxin pathway gene clusters in *Aspergillus parasiticus* and *Aspergillus flavus*. Applied and environmental microbiology, 61, 2365–2371.

12. Soldatou, S., Eldjarn, G.H., Huerta-Uribe, A., Rogers, S. and Duncan, K.R. (2019) Linking biosynthetic and chemical space to accelerate microbial secondary metabolite discovery. FEMS microbiology letters, 366.

13. Craney, A., Ahmed, S. and Nodwell, J. (2013) Towards a new science of secondary metabolism. The Journal of antibiotics, 66, 387–400.

14. Osbourn, A. (2010) Secondary metabolic gene clusters: evolutionary toolkits for chemical innovation. Trends in Genetics, 26, 449–457.

15. Keller, N.P. (2019) Fungal secondary metabolism: regulation, function and drug discovery. Nat Rev Microbiol, 17, 167–180.

16. Mulder, K.C., Mulinari, F., Franco, O.L., Soares, M.S., Magalhaes, B.S. and Parachin, N.S. (2015) Lovastatin production: From molecular basis to industrial process optimization. Biotechnol Adv, 33, 648–665.

17. Weber, G., Schörgendorfer, K., Schneider-Scherzer, E. and Leitner, E. (1994) The peptide synthetase catalyzing cyclosporine production in *Tolypocladium niveum* is encoded by a giant 45.8-kilobase open reading frame. Current Genetics, 26, 120–125.

18. van den Berg, M.A., Westerlaken, I., Leeflang, C., Kerkman, R. and Bovenberg, R.A. (2007) Functional characterization of the penicillin biosynthetic gene cluster of *Penicillium chrysogenum* Wisconsin54-1255. Fungal Genet Biol, 44, 830–844.

19. Nosanchuk, J.D., Stark, R.E. and Casadevall, A. (2015) Fungal Melanin: What do We Know About Structure? Front Microbiol, 6, 1463.

20. Wheeler, M.H. and Bell, A.A. (1988) Melanins and their importance in pathogenic fungi. Curr Top Med Mycol, 2, 338–387.

21. Luo, S. and Dong, S.H. (2019) Recent Advances in the Discovery and Biosynthetic Study of Eukaryotic RiPP Natural Products. Molecules, 24.

22. Montalban-Lopez, M., Scott, T.A., Ramesh, S., Rahman, I.R., van Heel, A.J., Viel, J.H., Bandarian, V., Dittmann, E., Genilloud, O., Goto, Y. et al. (2020) New developments in RiPP discovery, enzymology and engineering. Nat Prod Rep.

23. Blin, K., Shaw, S., Steinke, K., Villebro, R., Ziemert, N., Lee, S.Y., Medema, M.H. and Weber, T. (2019) antiSMASH 5.0: updates to the secondary metabolite genome mining pipeline. Nucleic Acids Research, 47, W81–W87.

24. Wolf, T., Shelest, V., Nath, N. and Shelest, E. (2016) CASSIS and SMIPS: promoter-based prediction of secondary metabolite gene clusters in eukaryotic genomes. Bioinformatics (Oxford, England), 32, 1138–1143.

25. Khaldi, N., Seifuddin, F.T., Turner, G., Haft, D., Nierman, W.C., Wolfe, K.H. and Fedorova, N.D. (2010) SMURF: Genomic mapping of fungal secondary metabolite clusters. Fungal genetics and biology: FG & B, 47, 736–741.

26. Hannigan, G.D., Prihoda, D., Palicka, A., Soukup, J., Klempir, O., Rampula, L., Durcak, J., Wurst, M., Kotowski, J., Chang, D. et al. (2019) A deep learning genome-mining strategy for biosynthetic gene cluster prediction. Nucleic acids research, 47, e110.

27. Sélem-Mojica, N., Aguilar, C., Gutiérrez-García, K., Martínez-Guerrero, C.E. and Barona-Gómez, F. (2019) EvoMining reveals the origin and fate of natural product biosynthetic enzymes. Microb Genom, 5.

28. Tai, Y., Liu, C., Yu, S., Yang, H., Sun, J., Guo, C., Huang, B., Liu, Z., Yuan, Y., Xia, E. et al. (2018) Gene co-expression network analysis reveals coordinated regulation of three characteristic secondary biosynthetic pathways in tea plant *(Camellia sinensis)*. BMC Genomics, 19, 616.

29. Ochoa, D. and Pazos, F. (2014) Practical aspects of protein co-evolution. Front Cell Dev Biol, 2, 14–14.

30. Stamatakis, A. (2014) RAxML version 8: a tool for phylogenetic analysis and post-analysis of large phylogenies. Bioinformatics, 30, 1312–1313.

31. Rice, P., Longden, I. and Bleasby, A. (2000) EMBOSS: the European Molecular Biology Open Software Suite. Trends Genet, 16, 276–277.

32. Altschul, S.F., Madden, T.L., Schäffer, A.A., Zhang, J., Zhang, Z., Miller, W. and Lipman, D.J. (1997) Gapped BLAST and PSI-BLAST: a new generation of protein database search programs. Nucleic Acids Res, 25, 3389–3402.

33. Thompson, J.D., Higgins, D.G. and Gibson, T.J. (1994) CLUSTAL W: improving the sensitivity of progressive multiple sequence alignment through sequence weighting, positions-specific gap penalties and weight matrix choice. Nucleic Acids Res, 22, 4673–4680.

34. Marcet-Houben, M. and Gabaldon, T. (2011) TreeKO: a duplication-aware algorithm for the comparison of phylogenetic trees. Nucleic Acids Res, 39, e66.

35. R Core Team. (2019) R: A language and environment for statistical computing. R Foundation for Statistical Computing.

36. Kucheryavskiy, S. (2019) mdatools: Multivariate Data Analysis for Chemometrics.

37. Warnes, G.R., Bolker, B., Bonebakker, L., Gentleman, R., Huber, W., Liaw, A., Lumley, T., Maechler, M., Magnusson, A., Moeller, S. et al. (2019) gplots: Various R Programming Tools for Plotting Data.

38. Wickham, H., Hester, J. and Francois, R. (2018) readr: Read Rectangular Text Data.

39. Fox, J. and Weisberg, S. (2019) An {R} Companion to Applied Regression. Sage.

40. Murtagh, F. and Legendre, P. (2014) Ward’s Hierarchical Agglomerative Clustering Method: Which Algorithms Implement Ward’s Criterion? Journal of Classification, 31, 274–295.

41. Nordberg, H., Cantor, M., Dusheyko, S., Hua, S., Poliakov, A., Shabalov, I., Smirnova, T., Grigoriev, I.V. and Dubchak, I. (2014) The genome portal of the Department of Energy Joint Genome Institute: 2014 updates. Nucleic Acids Res, 42, D26–31.

42. Terfehr, D., Dahlmann, T.A., Specht, T., Zadra, I., Kürnsteiner, H. and Kück, U. (2014) Genome Sequence and Annotation of *Acremonium chrysogenum*, Producer of the β-Lactam Antibiotic Cephalosporin C. Genome Announc, 2.

43. Nguyen, H.D., Lewis, C.T., Lévesque, C.A. and Gräfenhan, T. (2016) Draft Genome Sequence of *Alternaria alternata* ATCC 34957. Genome Announc, 4.

44. Hu, J., Chen, C., Peever, T., Dang, H., Lawrence, C. and Mitchell, T. (2012) Genomic characterization of the conditionally dispensable chromosome in *Alternaria arborescens* provides evidence for horizontal gene transfer. BMC Genomics, 13, 171.

45. Armitage, A.D., Cockerton, H.M., Sreenivasaprasad, S., Woodhall, J., Lane, C.R., Harrison, R.J. and Clarkson, J.P. (2020) Genomics Evolutionary History and Diagnostics of the *Alternaria alternata* Species Group Including Apple and Asian Pear Pathotypes. Frontiers in Microbiology, 10.

46. Lu, Y., Ye, C., Che, J., Xu, X., Shao, D., Jiang, C., Liu, Y. and Shi, J. (2019) Genomic sequencing, genome-scale metabolic network reconstruction, and in silico flux analysis of the grape endophytic fungus *Alternaria* sp. MG1. Microb Cell Fact, 18, 13.

47. Kohler, A., Kuo, A., Nagy, L.G., Morin, E., Barry, K.W., Buscot, F., Canbäck, B., Choi, C., Cichocki, N., Clum, A. et al. (2015) Convergent losses of decay mechanisms and rapid turnover of symbiosis genes in mycorrhizal mutualists. Nat Genet, 47, 410–415.

48. Martino, E., Morin, E., Grelet, G.A., Kuo, A., Kohler, A., Daghino, S., Barry, K.W., Cichocki, N., Clum, A., Dockter, R.B. et al. (2018) Comparative genomics and transcriptomics depict ericoid mycorrhizal fungi as versatile saprotrophs and plant mutualists. New Phytol, 217, 1213–1229.

49. Yang, J., Wang, L., Ji, X., Feng, Y., Li, X., Zou, C., Xu, J., Ren, Y., Mi, Q., Wu, J. et al. (2011) Genomic and proteomic analyses of the fungus *Arthrobotrys oligospora* provide insights into nematode-trap formation. PLoS Pathog, 7, e1002179.

50. Burmester, A., Shelest, E., Glöckner, G., Heddergott, C., Schindler, S., Staib, P., Heidel, A., Felder, M., Petzold, A., Szafranski, K. et al. (2011) Comparative and functional genomics provide insights into the pathogenicity of dermatophytic fungi. Genome Biol, 12, R7.

51. Murat, C., Payen, T., Noel, B., Kuo, A., Morin, E., Chen, J., Kohler, A., Krizsán, K., Balestrini, R., Da Silva, C. et al. (2018) *Pezizomycetes* genomes reveal the molecular basis of ectomycorrhizal truffle lifestyle. Nat Ecol Evol, 2, 1956–1965.

52. de Vries, R.P., Riley, R., Wiebenga, A., Aguilar-Osorio, G., Amillis, S., Uchima, C.A., Anderluh, G., Asadollahi, M., Askin, M., Barry, K. et al. (2017) Comparative genomics reveals high biological diversity and specific adaptations in the industrially and medically important fungal genus *Aspergillus*. Genome Biology, 18, 28.

53. Nierman, W.C., Yu, J., Fedorova-Abrams, N.D., Losada, L., Cleveland, T.E., Bhatnagar, D., Bennett, J. W., Dean, R. and Payne, G.A. (2015) Genome Sequence of *Aspergillus flavus* NRRL 3357, a Strain That Causes Aflatoxin Contamination of Food and Feed. Genome announcements, 3.

54. Nierman, W.C., Pain, A., Anderson, M.J., Wortman, J.R., Kim, H.S., Arroyo, J., Berriman, M., Abe, K., Archer, D.B., Bermejo, C. et al. (2005) Genomic sequence of the pathogenic and allergenic filamentous fungus *Aspergillus fumigatus*. Nature, 438, 1151–1156.

55. Pel, H.J., de Winde, J.H., Archer, D.B., Dyer, P.S., Hofmann, G., Schaap, P.J., Turner, G., de Vries, R.P., Albang, R., Albermann, K. et al. (2007) Genome sequencing and analysis of the versatile cell factory *Aspergillus niger* CBS 513.88. Nature biotechnology, 25, 221–231.

56. Zhao, G., Yao, Y., Qi, W., Wang, C., Hou, L., Zeng, B. and Cao, X. (2012) Draft genome sequence of *Aspergillus oryzae* strain 3.042. Eukaryot Cell, 11, 1178.

57. Vesth, T.C., Nybo, J.L., Theobald, S., Frisvad, J.C., Larsen, T.O., Nielsen, K.F., Hoof, J.B., Brandl, J., Salamov, A., Riley, R. et al. (2018) Investigation of inter- and intraspecies variation through genome sequencing of *Aspergillus* section Nigri. Nature Genetics, 50, 1688–1695.

58. Savitha, J., Bhargavi, S.D. and Praveen, V.K. (2016) Complete Genome Sequence of Soil Fungus *Aspergillus terreus* (KM017963), a Potent Lovastatin Producer. Genome Announc, 4.

59. Spanu, P.D., Abbott, J.C., Amselem, J., Burgis, T.A., Soanes, D.M., Stüber, K., Ver Loren van Themaat, E., Brown, J.K., Butcher, S.A., Gurr, S.J. et al. (2010) Genome expansion and gene loss in powdery mildew fungi reveal tradeoffs in extreme parasitism. Science, 330, 1543–1546.

60. Marsberg, A., Kemler, M., Jami, F., Nagel, J.H., Postma-Smidt, A., Naidoo, S., Wingfield, M.J., Crous, P.W., Spatafora, J.W., Hesse, C.N. et al. (2017) *Botryosphaeria dothidea:* a latent pathogen of global importance to woody plant health. Mol Plant Pathol, 18, 477–488.

61. Van Kan, J.A., Stassen, J.H., Mosbach, A., Van Der Lee, T.A., Faino, L., Farmer, A.D., Papasotiriou, D.G., Zhou, S., Seidl, M.F., Cottam, E. et al. (2017) A gapless genome sequence of the fungus *Botrytis cinerea*. Mol Plant Pathol, 18, 75–89.

62. Valero-Jiménez, C.A., Veloso, J., Staats, M. and van Kan, J.A.L. (2019) Comparative genomics of plant pathogenic *Botrytis* species with distinct host specificity. BMC Genomics, 20, 203.

63. Knapp, D.G., Németh, J.B., Barry, K., Hainaut, M., Henrissat, B., Johnson, J., Kuo, A., Lim, J.H.P., Lipzen, A., Nolan, M. et al. (2018) Comparative genomics provides insights into the lifestyle and reveals functional heterogeneity of dark septate endophytic fungi. Sci Rep, 8, 6321.

64. Teixeira, M.M., Moreno, L.F., Stielow, B.J., Muszewska, A., Hainaut, M., Gonzaga, L., Abouelleil, A., Patané, J.S., Priest, M., Souza, R. et al. (2017) Exploring the genomic diversity of black yeasts and relatives (Chaetothyriales, Ascomycota). Stud Mycol, 86, 1–28.

65. Cuomo, C.A., Untereiner, W.A., Ma, L.J., Grabherr, M. and Birren, B.W. (2015) Draft Genome Sequence of the Cellulolytic Fungus *Chaetomium globosum*. Genome announcements, 3.

66. Armaleo, D., Müller, O., Lutzoni, F., Andrésson Ó, S., Blanc, G., Bode, H.B., Collart, F.R., Dal Grande, F., Dietrich, F., Grigoriev, I.V. et al. (2019) The lichen symbiosis re-viewed through the genomes of *Cladonia grayi* and its algal partner *Asterochloris glomerata*. BMC Genomics, 20, 605.

67. Ohm, R.A., Feau, N., Henrissat, B., Schoch, C.L., Horwitz, B.A., Barry, K.W., Condon, B.J., Copeland, A.C., Dhillon, B., Glaser, F. et al. (2012) Diverse lifestyles and strategies of plant pathogenesis encoded in the genomes of eighteen *Dothideomycetes* fungi. PLoS pathogens, 8, e1003037.

68. Condon, B.J., Leng, Y., Wu, D., Bushley, K.E., Ohm, R.A., Otillar, R., Martin, J., Schackwitz, W., Grimwood, J., MohdZainudin, N. et al. (2013) Comparative genome structure, secondary metabolite, and effector coding capacity across *Cochliobolus* pathogens. PLoS Genet, 9, e1003233.

69. Baroncelli, R., Amby, D.B., Zapparata, A., Sarrocco, S., Vannacci, G., Le Floch, G., Harrison, R.J., Holub, E., Sukno, S.A., Sreenivasaprasad, S. et al. (2016) Gene family expansions and contractions are associated with host range in plant pathogens of the genus *Colletotrichum*. BMC Genomics, 17, 555.

70. Baroncelli, R., Sukno, S.A., Sarrocco, S., Cafà, G., Le Floch, G. and Thon, M.R. (2018) Whole-Genome Sequence of the Orchid Anthracnose Pathogen *Colletotrichum orchidophilum*. Mol Plant Microbe Interact, 31, 979–981.

71. Hacquard, S., Kracher, B., Hiruma, K., Münch, P.C., Garrido-Oter, R., Thon, M.R., Weimann, A., Damm, U., Dallery, J.F., Hainaut, M. et al. (2016) Survival trade-offs in plant roots during colonization by closely related beneficial and pathogenic fungi. Nat Commun, 7, 11362.

72. Lopez, D., Ribeiro, S., Label, P., Fumanal, B., Venisse, J.S., Kohler, A., de Oliveira, R.R., Labutti, K., Lipzen, A., Lail, K. et al. (2018) Genome-Wide Analysis of *Corynespora cassiicola* Leaf Fall Disease Putative Effectors. Front Microbiol, 9, 276.

73. Wu, W., Davis, R.W., Tran-Gyamfi, M.B., Kuo, A., LaButti, K., Mihaltcheva, S., Hundley, H., Chovatia, M., Lindquist, E., Barry, K. et al. (2017) Characterization of four endophytic fungi as potential consolidated bioprocessing hosts for conversion of lignocellulose into advanced biofuels. Appl Microbiol Biotechnol, 101, 2603–2618.

74. Morales-Cruz, A., Amrine, K.C., Blanco-Ulate, B., Lawrence, D.P., Travadon, R., Rolshausen, P.E., Baumgartner, K. and Cantu, D. (2015) Distinctive expansion of gene families associated with plant cell wall degradation, secondary metabolism, and nutrient uptake in the genomes of grapevine trunk pathogens. BMC Genomics, 16, 469.

75. Jones, L., Riaz, S., Morales-Cruz, A., Amrine, K.C., McGuire, B., Gubler, W.D., Walker, M.A. and Cantu, D. (2014) Adaptive genomic structural variation in the grape powdery mildew pathogen, *Erysiphe necator*. BMC Genomics, 15, 1081.

76. Blanco-Ulate, B., Rolshausen, P.E. and Cantu, D. (2013) Draft Genome Sequence of the Grapevine Dieback Fungus *Eutypa lata* UCR-EL1. Genome Announc, 1.

77. Bombassaro, A., de Hoog, S., Weiss, V.A., Souza, E.M., Leão, A.C., Costa, F.F., Baura, V., Tadra-Sfeir, M.Z., Balsanelli, E., Moreno, L.F. et al. (2016) Draft Genome Sequence of *Fonsecaea monophora* Strain CBS 269.37, an Agent of Human Chromoblastomycosis. Genome Announc, 4.

78. Bashyal, B.M., Rawat, K., Sharma, S., Kulshreshtha, D., Gopala Krishnan, S., Singh, A.K., Dubey, H., Solanke, A.U., Sharma, T.R. and Aggarwal, R. (2017) Whole Genome Sequencing of *Fusarium fujikuroi* Provides Insight into the Role of Secretory Proteins and Cell Wall Degrading Enzymes in Causing Bakanae Disease of Rice. Front Plant Sci, 8, 2013.

79. Cuomo, C.A., Guldener, U., Xu, J.R., Trail, F., Turgeon, B.G., Di Pietro, A., Walton, J.D., Ma, L.J., Baker, S.E., Rep, M. et al. (2007) The *Fusarium graminearum* genome reveals a link between localized polymorphism and pathogen specialization. Science, 317, 1400–1402.

80. Ma, L.J., van der Does, H.C., Borkovich, K.A., Coleman, J.J., Daboussi, M.J., Di Pietro, A., Dufresne, M., Freitag, M., Grabherr, M., Henrissat, B. et al. (2010) Comparative genomics reveals mobile pathogenicity chromosomes in *Fusarium*. Nature, 464, 367–373.

81. Niehaus, E.M., Münsterkötter, M., Proctor, R.H., Brown, D.W., Sharon, A., Idan, Y., Oren-Young, L., Sieber, C.M., Novák, O., Pěnčík, A. et al. (2016) Comparative “Omics” of the *Fusarium fujikuroi* Species Complex Highlights Differences in Genetic Potential and Metabolite Synthesis. Genome Biol Evol, 8, 3574–3599.

82. Gardiner, D.M., Benfield, A.H., Stiller, J., Stephen, S., Aitken, K., Liu, C. and Kazan, K. (2018) A high-resolution genetic map of the cereal crown rot pathogen *Fusarium pseudograminearum* provides a near-complete genome assembly. Mol Plant Pathol, 19, 217–226.

83. Okagaki, L.H., Nunes, C.C., Sailsbery, J., Clay, B., Brown, D., John, T., Oh, Y., Young, N., Fitzgerald, M., Haas, B.J. et al. (2015) Genome Sequences of Three Phytopathogenic Species of the *Magnaporthaceae* Family of Fungi. G3 (Bethesda), 5, 2539–2545.

84. Peter, M., Kohler, A., Ohm, R.A., Kuo, A., Krützmann, J., Morin, E., Arend, M., Barry, K.W., Binder, M., Choi, C. et al. (2016) Ectomycorrhizal ecology is imprinted in the genome of the dominant symbiotic fungus *Cenococcum geophilum*. Nat Commun, 7, 12662.

85. Dean, R.A., Talbot, N.J., Ebbole, D.J., Farman, M.L., Mitchell, T.K., Orbach, M.J., Thon, M., Kulkarni, R., Xu, J.R., Pan, H. et al. (2005) The genome sequence of the rice blast fungus *Magnaporthe grisea*. Nature, 434, 980–986.

86. Gao, Q., Jin, K., Ying, S.H., Zhang, Y., Xiao, G., Shang, Y., Duan, Z., Hu, X., Xie, X.Q., Zhou, G. et al. (2011) Genome sequencing and comparative transcriptomics of the model entomopathogenic fungi *Metarhizium anisopliae* and *M. acridum*. PLoS Genet, 7, e1001264.

87. Hu, X., Xiao, G., Zheng, P., Shang, Y., Su, Y., Zhang, X., Liu, X., Zhan, S., St Leger, R.J. and Wang, C. (2014) Trajectory and genomic determinants of fungal-pathogen speciation and host adaptation. Proc Natl Acad Sci U S A, 111, 16796–16801.

88. Binneck, E., Lastra, C.C.L. and Sosa-Gómez, D.R. (2019) Genome Sequence of *Metarhizium rileyi*, a Microbial Control Agent for *Lepidoptera*. Microbiol Resour Announc, 8.

89. Meerupati, T., Andersson, K.M., Friman, E., Kumar, D., Tunlid, A. and Ahrén, D. (2013) Genomic mechanisms accounting for the adaptation to parasitism in nematode-trapping fungi. PLoS Genet, 9, e1003909.

90. Tan, H., Kohler, A., Miao, R., Liu, T., Zhang, Q., Zhang, B., Jiang, L., Wang, Y., Xie, L., Tang, J. et al. (2019) Multi-omic analyses of exogenous nutrient bag decomposition by the black morel *Morchella importuna* reveal sustained carbon acquisition and transferring. Environ Microbiol, 21, 3909–3926.

91. Coleman, J.J., Rounsley, S.D., Rodriguez-Carres, M., Kuo, A., Wasmann, C.C., Grimwood, J., Schmutz, J., Taga, M., White, G.J., Zhou, S. et al. (2009) The genome of *Nectria haematococca*: contribution of supernumerary chromosomes to gene expansion. PLoS Genet, 5, e1000618.

92. Galagan, J.E., Calvo, S.E., Borkovich, K.A., Selker, E.U., Read, N.D., Jaffe, D., FitzHugh, W., Ma, L.J., Smirnov, S., Purcell, S. et al. (2003) The genome sequence of the filamentous fungus *Neurospora crassa*. Nature, 422, 859–868.

93. Baker, S.E., Schackwitz, W., Lipzen, A., Martin, J., Haridas, S., LaButti, K., Grigoriev, I.V., Simmons, B.A. and McCluskey, K. (2015) Draft Genome Sequence of *Neurospora crassa* Strain FGSC 73. Genome Announc, 3.

94. Ellison, C.E., Stajich, J.E., Jacobson, D.J., Natvig, D.O., Lapidus, A., Foster, B., Aerts, A., Riley, R., Lindquist, E.A., Grigoriev, I.V. et al. (2011) Massive changes in genome architecture accompany the transition to self-fertility in the filamentous fungus *Neurospora tetrasperma*. Genetics, 189, 55–69.

95. Haridas, S., Wang, Y., Lim, L., Massoumi Alamouti, S., Jackman, S., Docking, R., Robertson, G., Birol, I., Bohlmann, J. and Breuil, C. (2013) The genome and transcriptome of the pine saprophyte *Ophiostoma piceae*, and a comparison with the bark beetle-associated pine pathogen *Grosmannia clavigera*. BMC Genomics, 14, 373.

96. Urquhart, A.S., Mondo, S.J., Mäkelä, M.R., Hane, J.K., Wiebenga, A., He, G., Mihaltcheva, S., Pangilinan, J., Lipzen, A., Barry, K. et al. (2018) Genomic and Genetic Insights Into a Cosmopolitan Fungus, *Paecilomyces variotii (Eurotiales)*. Front Microbiol, 9, 3058.

97. Reynolds, H.T., Vijayakumar, V., Gluck-Thaler, E., Korotkin, H.B., Matheny, P.B. and Slot, J.C. (2018) Horizontal gene cluster transfer increased hallucinogenic mushroom diversity. Evol Lett, 2, 88–101.

98. Desjardins, C.A., Champion, M.D., Holder, J.W., Muszewska, A., Goldberg, J., Bailão, A.M., Brigido, M.M., Ferreira, M.E., Garcia, A.M., Grynberg, M. et al. (2011) Comparative genomic analysis of human fungal pathogens causing paracoccidioidomycosis. PLoS Genet, 7, e1002345.

99. Cheeseman, K., Ropars, J., Renault, P., Dupont, J., Gouzy, J., Branca, A., Abraham, A.L., Ceppi, M., Conseiller, E., Debuchy, R. et al. (2014) Multiple recent horizontal transfers of a large genomic region in cheese making fungi. Nat Commun, 5, 2876.

100. Specht, T., Dahlmann, T.A., Zadra, I., Kürnsteiner, H. and Kück, U. (2014) Complete Sequencing and Chromosome-Scale Genome Assembly of the Industrial Progenitor Strain P2niaD18 from the Penicillin Producer *Penicillium chrysogenum*. Genome Announc, 2.

101. Marcet-Houben, M., Ballester, A.R., de la Fuente, B., Harries, E., Marcos, J.F., González-Candelas, L. and Gabaldón, T. (2012) Genome sequence of the necrotrophic fungus *Penicillium digitatum*, the main postharvest pathogen of citrus. BMC Genomics, 13, 646.

102. Ballester, A.R., Marcet-Houben, M., Levin, E., Sela, N., Selma-Lázaro, C., Carmona, L., Wisniewski, M., Droby, S., González-Candelas, L. and Gabaldón, T. (2015) Genome, Transcriptome, and Functional Analyses of *Penicillium expansum* Provide New Insights Into Secondary Metabolism and Pathogenicity. Mol Plant Microbe Interact, 28, 232–248.

103. Nielsen, J.C., Grijseels, S., Prigent, S., Ji, B., Dainat, J., Nielsen, K.F., Frisvad, J.C., Workman, M. and Nielsen, J. (2017) Global analysis of biosynthetic gene clusters reveals vast potential of secondary metabolite production in *Penicillium species*. Nat Microbiol, 2, 17044.

104. Liu, G., Zhang, L., Wei, X., Zou, G., Qin, Y., Ma, L., Li, J., Zheng, H., Wang, S., Wang, C. et al. (2013) Genomic and secretomic analyses reveal unique features of the lignocellulolytic enzyme system of *Penicillium decumbens*. PLoS One, 8, e55185.

105. van den Berg, M.A., Albang, R., Albermann, K., Badger, J.H., Daran, J.M., Driessen, A.J., Garcia-Estrada, C., Fedorova, N.D., Harris, D.M., Heijne, W.H. et al. (2008) Genome sequencing and analysis of the filamentous fungus *Penicillium chrysogenum*. Nature biotechnology, 26, 1161–1168.

106. Wang, X., Zhang, X., Liu, L., Xiang, M., Wang, W., Sun, X., Che, Y., Guo, L., Liu, G., Guo, L. et al. (2015) Genomic and transcriptomic analysis of the endophytic fungus *Pestalotiopsis fici* reveals its lifestyle and high potential for synthesis of natural products. BMC Genomics, 16, 28.

107. Blanco-Ulate, B., Rolshausen, P. and Cantu, D. (2013) Draft Genome Sequence of the Ascomycete *Phaeoacremonium aleophilum* Strain UCR-PA7, a Causal Agent of the Esca Disease Complex in Grapevines. Genome Announc, 1.

108. Walker, A.K., Frasz, S.L., Seifert, K.A., Miller, J.D., Mondo, S.J., LaButti, K., Lipzen, A., Dockter, R.B., Kennedy, M.C., Grigoriev, I.V. et al. (2016) Full Genome of *Phialocephala scopiformis* DAOMC 229536, a Fungal Endophyte of Spruce Producing the Potent Anti-Insectan Compound Rugulosin. Genome Announc, 4.

109. Cissé, O.H., Pagni, M. and Hauser, P.M. (2012) *De novo* assembly of the *Pneumocystis jirovecii* genome from a single bronchoalveolar lavage fluid specimen from a patient. mBio, 4, e00428–00412.

110. Chibucos, M.C., Crabtree, J., Nagaraj, S., Chaturvedi, S. and Chaturvedi, V. (2013) Draft Genome Sequences of Human Pathogenic Fungus *Geomyces pannorum* Sensu Lato and Bat White Nose Syndrome Pathogen *Geomyces (Pseudogymnoascus) destructans*. Genome Announc, 1.

111. Mondo, S.J., Dannebaum, R.O., Kuo, R.C., Louie, K.B., Bewick, A.J., LaButti, K., Haridas, S., Kuo, A., Salamov, A., Ahrendt, S.R. et al. (2017) Widespread adenine N6-methylation of active genes in fungi. Nat Genet, 49, 964–968.

112. Cubeta, M.A., Thomas, E., Dean, R.A., Jabaji, S., Neate, S.M., Tavantzis, S., Toda, T., Vilgalys, R., Bharathan, N., Fedorova-Abrams, N. et al. (2014) Draft Genome Sequence of the Plant-Pathogenic Soil Fungus *Rhizoctonia solani* Anastomosis Group 3 Strain Rhs1AP. Genome Announc, 2.

113. Liti, G., Nguyen Ba, A.N., Blythe, M., Müller, C.A., Bergström, A., Cubillos, F.A., Dafhnis-Calas, F., Khoshraftar, S., Malla, S., Mehta, N. et al. (2013) High quality *de novo* sequencing and assembly of the *Saccharomyces arboricolus* genome. BMC Genomics, 14, 69.

114. Goffeau, A., Barrell, B.G., Bussey, H., Davis, R.W., Dujon, B., Feldmann, H., Galibert, F., Hoheisel, J.D., Jacq, C., Johnston, M. et al. (1996) Life with 6000 genes. Science, 274, 546, 563–547.

115. Quandt, C.A., Bushley, K.E. and Spatafora, J.W. (2015) The genome of the truffle-parasite *Tolypocladium ophioglossoides* and the evolution of antifungal peptaibiotics. BMC Genomics, 16, 553.

116. Quandt, C.A., Patterson, W. and Spatafora, J.W. (2018) Harnessing the power of phylogenomics to disentangle the directionality and signatures of interkingdom host jumping in the parasitic fungal genus *Tolypocladium*. Mycologia, 110, 104–117.

117. Proctor, R.H., McCormick, S.P., Kim, H.S., Cardoza, R.E., Stanley, A.M., Lindo, L., Kelly, A., Brown, D.W., Lee, T., Vaughan, M.M. et al. (2018) Evolution of structural diversity of trichothecenes, a family of toxins produced by plant pathogenic and entomopathogenic fungi. PLoS Pathog, 14, e1006946.

118. Druzhinina, I.S., Chenthamara, K., Zhang, J., Atanasova, L., Yang, D., Miao, Y., Rahimi, M.J., Grujic, M., Cai, F., Pourmehdi, S. et al. (2018) Massive lateral transfer of genes encoding plant cell wall-degrading enzymes to the mycoparasitic fungus *Trichoderma* from its plant-associated hosts. PLoS Genet, 14, e1007322.

119. Kubicek, C.P., Herrera-Estrella, A., Seidl-Seiboth, V., Martinez, D.A., Druzhinina, I.S., Thon, M., Zeilinger, S., Casas-Flores, S., Horwitz, B.A., Mukherjee, P.K. et al. (2011) Comparative genome sequence analysis underscores mycoparasitism as the ancestral life style of *Trichoderma*. Genome Biol, 12, R40.

120. Baroncelli, R., Piaggeschi, G., Fiorini, L., Bertolini, E., Zapparata, A., Pè, M.E., Sarrocco, S. and Vannacci, G. (2015) Draft Whole-Genome Sequence of the Biocontrol Agent *Trichoderma harzianum* T6776. Genome Announc, 3.

121. Xie, B.B., Qin, Q.L., Shi, M., Chen, L.L., Shu, Y.L., Luo, Y., Wang, X.W., Rong, J.C., Gong, Z.T., Li, D. et al. (2014) Comparative genomics provide insights into evolution of *trichoderma* nutrition style. Genome Biol Evol, 6, 379–390.

122. Martinez, D., Berka, R.M., Henrissat, B., Saloheimo, M., Arvas, M., Baker, S.E., Chapman, J., Chertkov, O., Coutinho, P.M., Cullen, D. et al. (2008) Genome sequencing and analysis of the biomass-degrading fungus *Trichoderma reesei* (syn. *Hypocrea jecorina*). Nature biotechnology, 26, 553–560.

123. Martinez, D.A., Oliver, B.G., Gräser, Y., Goldberg, J.M., Li, W., Martinez-Rossi, N.M., Monod, M., Shelest, E., Barton, R.C., Birch, E. et al. (2012) Comparative genome analysis of *Trichophyton rubrum* and related dermatophytes reveals candidate genes involved in infection. mBio, 3, e00259–00212.

124. Deng, C.H., Plummer, K.M., Jones, D.A.B., Mesarich, C.H., Shiller, J., Taranto, A.P., Robinson, A.J., Kastner, P., Hall, N.E., Templeton, M.D. et al. (2017) Comparative analysis of the predicted secretomes of *Rosaceae* scab pathogens *Venturia inaequalis* and *V. pirina* reveals expanded effector families and putative determinants of host range. BMC Genomics, 18, 339.

125. Klosterman, S.J., Subbarao, K.V., Kang, S., Veronese, P., Gold, S.E., Thomma, B.P., Chen, Z., Henrissat, B., Lee, Y.H., Park, J. et al. (2011) Comparative genomics yields insights into niche adaptation of plant vascular wilt pathogens. PLoS Pathog, 7, e1002137.

126. Gazis, R., Kuo, A., Riley, R., LaButti, K., Lipzen, A., Lin, J., Amirebrahimi, M., Hesse, C.N., Spatafora, J.W., Henrissat, B. et al. (2016) The genome of *Xylona heveae* provides a window into fungal endophytism. Fungal Biol, 120, 26–42.

127. Grandaubert, J., Bhattacharyya, A. and Stukenbrock, E.H. (2015) RNA-seq-Based Gene Annotation and Comparative Genomics of Four Fungal Grass Pathogens in the Genus *Zymoseptoria* Identify Novel Orphan Genes and Species-Specific Invasions of Transposable Elements. G3 (Bethesda), 5, 1323–1333.

128. Stukenbrock, E.H., Christiansen, F.B., Hansen, T.T., Dutheil, J.Y. and Schierup, M.H. (2012) Fusion of two divergent fungal individuals led to the recent emergence of a unique widespread pathogen species. Proc Natl Acad Sci U S A, 109, 10954–10959.

129. Bailey, T.L., Boden, M., Buske, F.A., Frith, M., Grant, C.E., Clementi, L., Ren, J., Li, W.W. and Noble, W.S. (2009) MEME SUITE: tools for motif discovery and searching. Nucleic acids research, 37, W202–208.

130. Kautsar, S.A., Blin, K., Shaw, S., Navarro-Muñoz, J.C., Terlouw, B.R., van der Hooft, J.J.J., van Santen, J.A., Tracanna, V., Suarez Duran, H.G., Pascal Andreu, V. et al. (2019) MIBiG 2.0: a repository for biosynthetic gene clusters of known function. Nucleic Acids Research, 48, D454–D458.

131. Heneghan, M.N., Yakasai, A.A., Williams, K., Kadir, K.A., Wasil, Z., Bakeer, W., Fisch, K.M., Bailey, A.M., Simpson, T.J., Cox, R.J. et al. (2011) The programming role of trans-acting enoyl reductases during the biosynthesis of highly reduced fungal polyketides. Chemical Science, 2.

132. Ehrlich, K.C., Chang, P.K., Yu, J. and Cotty, P.J. (2004) Aflatoxin biosynthesis cluster gene *cypA* is required for G aflatoxin formation. Appl Environ Microbiol, 70, 6518–6524.

133. Bhatnagar, D., Cary, J.W., Ehrlich, K., Yu, J. and Cleveland, T.E. (2006) Understanding the genetics of regulation of aflatoxin production and *Aspergillus flavus* development. Mycopathologia, 162, 155–166.

134. Porquier, A., Morgant, G., Moraga, J., Dalmais, B., Luyten, I., Simon, A., Pradier, J.M., Amselem, J., Collado, I.G. and Viaud, M. (2016) The botrydial biosynthetic gene cluster of *Botrytis cinerea* displays a bipartite genomic structure and is positively regulated by the putative Zn(II)2Cys6 transcription factor BcBot6. Fungal Genet Biol, 96, 33–46.

135. Pinedo, C., Wang, C.M., Pradier, J.M., Dalmais, B., Choquer, M., Le Pecheur, P., Morgant, G., Collado, I.G., Cane, D.E. and Viaud, M. (2008) Sesquiterpene synthase from the botrydial biosynthetic gene cluster of the phytopathogen *Botrytis cinerea*. ACS Chem Biol, 3, 791–801.

136. Hamed, R.B., Gomez-Castellanos, J.R., Henry, L., Ducho, C., McDonough, M.A. and Schofield, C.J. (2013) The enzymes of beta-lactam biosynthesis. Nat Prod Rep, 30, 21–107.

137. Abe, Y., Suzuki, T., Ono, C., Iwamoto, K., Hosobuchi, M. and Yoshikawa, H. (2002) Molecular cloning and characterization of an ML-236B (compactin) biosynthetic gene cluster in *Penicillium citrinum*. Mol Genet Genomics, 267, 636–646.

138. Abe, Y., Ono, C., Hosobuchi, M. and Yoshikawa, H. (2002) Functional analysis of *mlcR*, a regulatory gene for ML-236B (compactin) biosynthesis in *Penicillium citrinum*. Mol Genet Genomics, 268, 352–361.

139. Hoffmann, K., Schneider-Scherzer, E., Kleinkauf, H. and Zocher, R. (1994) Purification and Characterization of Eucaryotic Alanine Racemase Acting as Key Enzyme in Cyclosporin Biosynthesis. Biological Chemistry, 269, 12710–12714.

140. Yang, X., Feng, P., Yin, Y., Bushley, K., Spatafora, J.W. and Wang, C. (2018) Cyclosporine Biosynthesis in *Tolypocladium inflatum* Benefits Fungal Adaptation to the Environment. Molecular Biology and Physiology, 9.

141. Weber, G. and Leitner, E. (1994) Disruption of the cyclosporin synthetase gene of *Tolypocladium niveum*. Current Genetics, 26, 461 – 467.

142. Wang, B., Kang, Q., Lu, Y., Bai, L. and Wang, C. (2012) Unveiling the biosynthetic puzzle of destruxins in *Metarhizium* species. Proc Natl Acad Sci U S A, 109, 1287–1292.

143. Lin, H.C., Chooi, Y.H., Dhingra, S., Xu, W., Calvo, A.M. and Tang, Y. (2013) The fumagillin biosynthetic gene cluster in *Aspergillus fumigatus* encodes a cryptic terpene cyclase involved in the formation of beta-trans-bergamotene. J Am Chem Soc, 135, 4616–4619.

144. Kato, N., Suzuki, H., Takagi, H., Uramoto, M., Takahashi, S. and Osada, H. (2011) Gene disruption and biochemical characterization of verruculogen synthase of *Aspergillus fumigatus*. Chembiochem, 12, 711–714.

145. Kato, N., Suzuki, H., Takagi, H., Asami, Y., Kakeya, H., Uramoto, M., Usui, T., Takahashi, S., Sugimoto, Y. and Osada, H. (2009) Identification of cytochrome P450s required for fumitremorgin biosynthesis in *Aspergillus fumigatus*. Chembiochem, 10, 920–928.

146. Maiya, S., Grundmann, A., Li, S.M. and Turner, G. (2009) Improved tryprostatin B production by heterologous gene expression in *Aspergillus nidulans*. Fungal Genet Biol, 46, 436–440.

147. Kato, N., Suzuki, H., Okumura, H., Takahashi, S. and Osada, H. (2013) A point mutation in *ftmD* blocks the fumitremorgin biosynthetic pathway in *Aspergillus fumigatus* strain Af293. Biosci Biotechnol Biochem, 77, 1061–1067.

148. Proctor, R.H., Busman, M., Seo, J.A., Lee, Y.W. and Plattner, R.D. (2008) A fumonisin biosynthetic gene cluster in *Fusarium oxysporum* strain O-1890 and the genetic basis for B versus C fumonisin production. Fungal Genet Biol, 45, 1016–1026.

149. Zaleta-Rivera, K., Xu, C., Yu, F., Butchko, R.A., Proctor, R.H., Hidalgo-Lara, M.E., Raza, A., Dussault, P.H. and Du, L. (2006) A Bidomain Nonribosomal Peptide Synthetase Encoded by FUM14 Catalyzes the Formation of Tricarballylic Esters in the Biosynthesis of Fumonisins. Biochemistry 45, 2561–2569.

150. Butchko, R.A., Plattner, R.D. and Proctor, R.H. (2006) Deletion Analysis of FUM Genes Involved in Tricarballylic Ester Formation during Fumonisin Biosynthesis. J. Agric. Food Chem., 54, 9398–9404.

151. Butchko, R.A., Plattner, R.D. and Proctor, R.H. (2003) FUM13 Encodes a Short Chain Dehydrogenase/Reductase Required for C-3 Carbonyl Reduction during Fumonisin Biosynthesis in *Gibberella moniliformis*. J. Agric. Food Chem., 51, 3000–3006.

152. Butchko, R.A., Plattner, R.D. and Proctor, R.H. (2003) FUM9 is required for C-5 hydroxylation of fumonisins and complements the meitotically defined Fum3 locus in *Gibberella moniliformis*. Appl Environ Microbiol, 69, 6935–6937.

153. Brown, D.W., Butchko, R.A., Busman, M. and Proctor, R.H. (2007) The *Fusarium verticillioides* FUM gene cluster encodes a Zn(II)2Cys6 protein that affects FUM gene expression and fumonisin production. Eukaryot Cell, 6, 1210–1218.

154. Du, L., Zhu, X., Gerber, R., Huffman, J., Lou, L., Jorgenson, J., Yu, F., Zaleta-Rivera, K. and Wang, Q. (2008) Biosynthesis of sphinganine-analog mycotoxins. J Ind Microbiol Biotechnol, 35, 455–464.

155. Proctor, R.H., Desjardins, A.E., Plattner, R.D. and Hohn, T.M. (1999) A Polyketide Synthase Gene Required for Biosynthesis of Fumonisin Mycotoxins in *Gibberella fujikuroi* Mating Population A. Fungal Genetics and Biology 27, 100–112.

156. Lia, Y., Lou, L., Cerny, R.L., Butchko, R.A., Proctor, R.H., Shen, Y. and Du, L. (2013) Tricarballylic ester formation during biosynthesis of fumonisin mycotoxins in *Fusarium verticillioides*. Mycology, 4, 179–186.

157. Studt, L., Janevska, S., Niehaus, E.M., Burkhardt, I., Arndt, B., Sieber, C.M., Humpf, H.U., Dickschat, J.S. and Tudzynski, B. (2016) Two separate key enzymes and two pathway-specific transcription factors are involved in fusaric acid biosynthesis in *Fusarium fujikuroi*. Environ Microbiol, 18, 936–956.

158. Lin, X., Yuan, S., Chen, S., Chen, B., Xu, H., Liu, L., Li, H. and Gao, Z. (2019) Heterologous Expression of Ilicicolin H Biosynthetic Gene Cluster and Production of a New Potent Antifungal Reagent, Ilicicolin J. Molecules, 24.

159. Cary, J.W., Uka, V., Han, Z., Buyst, D., Harris-Coward, P.Y., Ehrlich, K.C., Wei, Q., Bhatnagar, D., Dowd, P.F., Martens, S.L. et al. (2015) An *Aspergillus flavus* secondary metabolic gene cluster containing a hybrid PKS-NRPS is necessary for synthesis of the 2-pyridones, leporins. Fungal Genet Biol, 81, 88–97.

160. Manzoni, M. and Rollini, M. (2002) Biosynthesis and biotechnological production of statins by filamentous fungi and application of these cholesterol-lowering drugs. Appl Microbiol Biotechnol, 58, 555–564.

161. Zhang, W., Cao, S., Qiu, L., Qi, F., Li, Z., Yang, Y., Huang, S., Bai, F., Liu, C., Wan, X. et al. (2015) Functional characterization of MpaG′, the O-methyltransferase involved in the biosynthesis of mycophenolic acid. Chembiochem, 16, 565–569.

162. Zhang, W., Du, L., Qu, Z., Zhang, X., Li, F., Li, Z., Qi, F., Wang, X., Jiang, Y., Men, P. et al. (2019) Compartmentalized biosynthesis of mycophenolic acid. Proc Natl Acad Sci U S A, 116, 13305–13310.

163. Hansen, B.G., Genee, H.J., Kaas, C.S., Nielsen, J.B., Regueira, T.B., Mortensen, U.H., Frisvad, J.C. and Patil, K.R. (2011) A new class of IMP dehydrogenase with a role in self-resistance of mycophenolic acid producing fungi. BMC Microbiol, 11, 202.

164. Hansen, B.G., Salomonsen, B., Nielsen, M.T., Nielsen, J.B., Hansen, N.B., Nielsen, K.F., Regueira, T.B., Nielsen, J., Patil, K.R. and Mortensen, U.H. (2011) Versatile enzyme expression and characterization system for *Aspergillus nidulans*, with the *Penicillium brevicompactum* polyketide synthase gene from the mycophenolic acid gene cluster as a test case. Appl Environ Microbiol, 77, 3044–3051.

165. Regueira, T.B., Kildegaard, K.R., Hansen, B.G., Mortensen, U.H., Hertweck, C. and Nielsen, J. (2011) Molecular basis for mycophenolic acid biosynthesis in *Penicillium brevicompactum*. Appl Environ Microbiol, 77, 3035–3043.

166. Hansen, B.G., Mnich, E., Nielsen, K.F., Nielsen, J.B., Nielsen, M.T., Mortensen, U.H., Larsen, T.O. and Patil, K.R. (2012) Involvement of a natural fusion of a cytochrome P450 and a hydrolase in mycophenolic acid biosynthesis. Appl Environ Microbiol, 78, 4908–4913.

167. Del-Cid, A., Gil-Duran, C., Vaca, I., Rojas-Aedo, J.F., Garcia-Rico, R.O., Levican, G. and Chavez, R. (2016) Identification and Functional Analysis of the Mycophenolic Acid Gene Cluster of *Penicillium roqueforti*. PLoS One, 11, e0147047.

168. Gillot, G., Jany, J.L., Dominguez-Santos, R., Poirier, E., Debaets, S., Hidalgo, P.I., Ullan, R.V., Coton, E. and Coton, M. (2017) Genetic basis for mycophenolic acid production and strain-dependent production variability in *Penicillium roqueforti*. Food Microbiol, 62, 239–250.

169. Scott, B., Young, C.A., Saikia, S., McMillan, L.K., Monahan, B.J., Koulman, A., Astin, J., Eaton, C.J., Bryant, A., Wrenn, R.E. et al. (2013) Deletion and gene expression analyses define the paxilline biosynthetic gene cluster in *Penicillium paxilli*. Toxins (Basel), 5, 1422–1446.

170. Fierro, F., Garcia-Estrada, C., Castillo, N.I., Rodriguez, R., Velasco-Conde, T. and Martin, J.F. (2006) Transcriptional and bioinformatic analysis of the 56.8 kb DNA region amplified in tandem repeats containing the penicillin gene cluster in *Penicillium chrysogenum*. Fungal Genet Biol, 43, 618–629.

171. Xu, X., Liu, L., Zhang, F., Wang, W., Li, J., Guo, L., Che, Y. and Liu, G. (2014) Identification of the first diphenyl ether gene cluster for pestheic acid biosynthesis in plant endophyte *Pestalotiopsis fici*. Chembiochem, 15, 284–292.

172. Chen, L., Yue, Q., Zhang, X., Xiang, M., Wang, C., Li, S., Che, Y., Ortiz-López, F.J., Bills, G.F., Liu, X. et al. (2013) Genomics-driven discovery of the pneumocandin biosynthetic gene cluster in the fungus *Glarea lozoyensis*. BMC Genomics, 14.

173. Chen, L., Li, Y., Yue, Q., Loksztejn, A., Yokoyama, K., Felix, E.A., Liu, X., Zhang, N., An, Z. and Bills, G.F. (2016) Engineering of New Pneumocandin Side-Chain Analogues from *Glarea lozoyensis* by Mutasynthesis and Evaluation of Their Antifungal Activity. ACS Chem Biol, 11, 2724–2733.

174. Chen, L., Yue, Q., Li, Y., Niu, X., Xiang, M., Wang, W., Bills, G.F., Liu, X. and An, Z. (2015) Engineering of *Glarea lozoyensis* for exclusive production of the pneumocandin B0 precursor of the antifungal drug caspofungin acetate. Appl Environ Microbiol, 81, 1550–1558.

175. Salo, O., Guzman-Chavez, F., Ries, M.I., Lankhorst, P.P., Bovenberg, R.A.L., Vreeken, R.J. and Driessen, A.J.M. (2016) Identification of a Polyketide Synthase Involved in Sorbicillin Biosynthesis by *Penicillium chrysogenum*. Appl Environ Microbiol, 82, 3971–3978.

176. Guzman-Chavez, F., Salo, O., Nygard, Y., Lankhorst, P.P., Bovenberg, R.A.L. and Driessen, A.J.M. (2017) Mechanism and regulation of sorbicillin biosynthesis by *Penicillium chrysogenum*. Microb Biotechnol, 10, 958–968.

177. Derntl, C., Guzman-Chavez, F., Mello-de-Sousa, T.M., Busse, H.J., Driessen, A.J.M., Mach, R.L. and Mach-Aigner, A.R. (2017) *In Vivo* Study of the Sorbicillinoid Gene Cluster in *Trichoderma reesei*. Frontiers in microbiology, 8, 2037.

178. Heneghan, M.N., Yakasai, A.A., Halo, L.M., Song, Z., Bailey, A.M., Simpson, T.J., Cox, R.J. and Lazarus, C.M. (2010) First heterologous reconstruction of a complete functional fungal biosynthetic multigene cluster. Chembiochem, 11, 1508–1512.

179. Halo, L.M., Heneghan, M.N., Yakasai, A.A., Song, Z., Williams, K., Bailey, A.M., Cox, R.J., Lazarus, C.M. and Simpson, T.J. (2008) Late Stage Oxidations during the Biosynthesis of the 2-Pyridone Tenellin in the Entomopathogenic Fungus *Beauveria bassiana*. J. Am. Chem. Soc., 130, 17988–17996.

180. Zaehle, C., Gressler, M., Shelest, E., Geib, E., Hertweck, C. and Brock, M. (2014) Terrein biosynthesis in *Aspergillus terreus* and its impact on phytotoxicity. Chem Biol, 21, 719–731.

181. Kakule, T.B., Zhang, S., Zhan, J. and Schmidt, E.W. (2015) Biosynthesis of the tetramic acids Sch210971 and Sch210972. Org Lett, 17, 2295–2297.

182. Umemura, M., Nagano, N., Koike, H., Kawano, J., Ishii, T., Miyamura, Y., Kikuchi, M., Tamano, K., Yu, J., Shin-ya, K. et al. (2014) Characterization of the biosynthetic gene cluster for the ribosomally synthesized cyclic peptide ustiloxin B in *Aspergillus flavus*. Fungal Genet Biol, 68, 23–30.

183. Lim, F.Y., Won, T.H., Raffa, N., Baccile, J.A., Wisecaver, J., Rokas, A., Schroeder, F.C. and Keller, N.P. (2018) Fungal Isocyanide Synthases and Xanthocillin Biosynthesis in *Aspergillus fumigatus*. mBio, 9.

184. Chicco, D. and Jurman, G. (2020) The advantages of the Matthews correlation coefficient (MCC) over F1 score and accuracy in binary classification evaluation. BMC Genomics, 21, 6.

185. Buchfink, B., Xie, C. and Huson, D.H. (2015) Fast and sensitive protein alignment using DIAMOND. Nat Methods, 12, 59–60.

186. Finn, R.D., Clements, J. and Eddy, S.R. (2011) HMMER web server: interactive sequence similarity searching. Nucleic Acids Research, 39, W29–W37.

