## Supplementary data file 2 for "The Functional Order (FunOrder) tool – Identification of essential biosynthetic genes through computational molecular co-evolution"

**Table S1.** Definition of topology

**Table S2.** Parameters used to calculate the manual evaluation measure (MEM)

**Table S3.** Confusion matrix of MEM values

**Table S4.** Negative control value (NCV) of the random BGCs calculated with equation 3 and classification as true negative (TN) or false positive (FP) based on the threshold of 0.66

**Table S5.** Strict distance matrix of the Lovastatin BGC of *Aspergillus terreus* (lov)

**Table S6.** Evolutionary distance matrix of the Lovastatin BGC of *Aspergillus terreus* (lov)

**Table S7.** Combined distance matrix of the Lovastatin BGC of *Aspergillus terreus* (lov)

**Figure S1.** Heatmap of the strict distance matrix of the Lovastatin BGC of *Aspergillus terreus* (lov).

**Figure S2.** Heatmap of the evolutionary distance matrix of the Lovastatin BGC of *Aspergillus terreus* (lov).

**Figure S3.** Heatmap of the combined distance matrix of the Lovastatin BGC of *Aspergillus terreus* (lov).

**Figure S4.** Dendrograms based on the scaled (A) and unscaled (B) strict distance matrix of the Lovastatin BGC of *Aspergillus terreus* (lov).

**Figure S5.** Dendrograms based on the scaled (A) and unscaled (B) evolutionary distance matrix of the Lovastatin BGC of *Aspergillus terreus* (lov).

**Figure S6.** Dendrograms based on the scaled (A) and unscaled (B) combined distance matrix of the Lovastatin BGC of *Aspergillus terreus* (lov).

**Figure S7.** Score plot of the first two principal components from a PCA performed on the evolutionary distance matrix analysis of the Lovastatin BGC of *Aspergillus terreus* (lov).

**Table S1.** Definition of topology

| Topology | Definition |
|---|---|
| **same** | min. 8 similar species, same topology with only little exceptions, colour 70-100% |
| **very similar** | min.5 similar species, similar topology, colour min. 70% |
| **similar** | distance < 2, colour min. 50% |
| **somewhat similar** | either 1 or 2 similar species with distances < 0.5 and nodes <3, or more species but only little similarities |
| **different** | no similarities or only 1 similar species |

**Table S2.** Parameters used to calculate the manual evaluation measure (MEM)

| ΔDistance | ΔNodes | Color | Topology | MEM |
|---|---|---|---|---|
| **0 – 0.5** | 0 | blue | same | 3 |
| **0.5 - 1** | 1 | - | very similar | 2.5 |
| **1 – 1.5** | 2 | green | Similar | 2 |
| **1.5 - 2** | 3 | - | somewhat similar | 1.5 |
| **> 2** | > 4 | yellow | different | 1 |

**Table S3.** Confusion matrix of MEM values

| MEM | Predicted | |
|---|---|---|
| | positive | negative |
| **positive** | 21 | 9 |
| **negative** | 8 | 52 |

**Table S4.** Negative control value (NCV) of the random BGCs calculated with equation 3 and classification as true negative (TN) or false positive (FP) based on the threshold of 0.66

| Random BGC | NCV (0-1) | TN / FP |
|---|---|---|
| **1** | 0.80 | TN |
| **2** | 0.66 | TN |
| **3** | 0.64 | FP |
| **4** | 0.83 | TN |
| **5** | 0.66 | TN |
| **6** | 1 | TN |
| **7** | 0.66 | TN |
| **8** | 1 | TN |
| **9** | 0.66 | TN |
| **10** | 1 | TN |
| **11** | 0.81 | TN |
| **12** | 0.75 | TN |
| **13** | 0.60 | FP |
| **14** | 0.66 | TN |
| **15** | 0.66 | TN |
| **16** | 1 | TN |
| **17** | 0.83 | TN |
| **18** | 0.81 | TN |
| **19** | 0.73 | TN |
| **20** | 0.70 | TN |
| **21** | 1 | TN |
| **22** | 0.95 | TN |
| **23** | 0.92 | TN |
| **24** | 0.62 | FP |
| **25** | 0.68 | TN |
| **26** | 0.86 | TN |
| **27** | 0.65 | FP |
| **28** | 0.77 | TN |
| **29** | 0.74 | TN |
| **30** | 0.50 | FP |
| **31** | 0.61 | FP |
| **32** | 0.83 | TN |
| **33** | 0.80 | TN |
| **34** | 0.75 | TN |
| **35** | 0.75 | TN |
| **36** | 0.75 | TN |
| **37** | 0.83 | TN |
| **38** | 0.66 | TN |
| **39** | 0.90 | TN |
| **40** | 0.76 | TN |
| **41** | 0.55 | FP |
| **42** | 0.83 | TN |
| **43** | 0.58 | FP |
| **44** | 0.76 | TN |
| **45** | 0.76 | TN |

**Table S4 (continued).** Negative control value (NCV) of the random BGCs calculated with equation 3 and classification as true negative (TN) or false positive (FP) based on the threshold of 0.66

| Random BGC | NCV (0-1) | TN / FP |
|---|---|---|
| **46** | 0.59 | FP |
| **47** | 0.75 | TN |
| **48** | 0.85 | TN |
| **49** | 0.50 | FP |
| **50** | 0.83 | TN |
| **51** | 0.76 | TN |
| **52** | 0.83 | TN |
| **53** | 0.70 | TN |
| **54** | 0.80 | TN |
| **55** | 0.65 | FP |
| **56** | 0.81 | TN |
| **57** | 0.73 | TN |
| **58** | 0.75 | TN |
| **59** | 0.70 | TN |
| **60** | 1 | TN |

**Table S5.** Strict distance matrix of the Lovastatin BGC of *Aspergillus terreus* (lov)

| | orf1 | orf2 | lovA | lovB | lovG | lovC | lovD | orf80c | lovE | orf10c | lovF | orf13 | orf14 | orf15 | orf16 | orf17 | orf18 |
|---|---|---|---|---|---|---|---|---|---|---|---|---|---|---|---|---|---|
| **orf1** | 0 | 0.853 | 0.915 | 0.857 | 0.9 | 0.891 | 0.932 | 0.909 | 0.806 | 0.944 | 0.899 | 0.884 | 0.899 | 0.883 | 0.918 | 0.912 | 0.804 |
| **orf2** | 0.853 | 0 | 0.873 | 0.824 | 0.864 | 0.845 | 0.904 | 0.675 | 0.811 | 0.695 | 0.807 | 0.806 | 0.951 | 0.629 | 0.83 | 0.705 | 0.773 |
| **lovA** | 0.915 | 0.873 | 0 | 0.829 | 0.711 | 0.918 | 0.738 | 0.873 | 0.844 | 0.725 | 0.603 | 0.839 | 0.924 | 0.859 | 0.96 | 0.857 | 0.791 |
| **lovB** | 0.857 | 0.824 | 0.829 | 0 | 0.57 | 0.66 | 0.746 | 0.87 | 0.763 | 0.805 | 0.771 | 0.904 | 0.827 | 0.916 | 0.81 | 0.829 | 0.888 |
| **lovG** | 0.9 | 0.864 | 0.711 | 0.57 | 0 | 0.813 | 0.845 | 0.907 | 0.717 | 0.734 | 0.708 | 0.916 | 0.901 | 0.889 | 0.915 | 0.922 | 0.727 |
| **lovC** | 0.891 | 0.845 | 0.918 | 0.66 | 0.813 | 0 | 0.863 | 0.834 | 0.808 | 0.866 | 0.86 | 0.862 | 0.945 | 0.882 | 0.848 | 0.852 | 1 |
| **lovD** | 0.932 | 0.904 | 0.738 | 0.746 | 0.845 | 0.863 | 0 | 0.968 | 1 | 0.824 | 0.824 | 0.949 | 1 | 0.931 | 0.97 | 0.944 | 0.871 |
| **orf80c** | 0.909 | 0.675 | 0.873 | 0.87 | 0.907 | 0.834 | 0.968 | 0 | 0.802 | 0.714 | 0.755 | 0.669 | 0.95 | 0.674 | 0.495 | 0.557 | 0.722 |
| **lovE** | 0.806 | 0.811 | 0.844 | 0.763 | 0.717 | 0.808 | 1 | 0.802 | 0 | 0.807 | 0.825 | 0.784 | 0.786 | 0.712 | 0.808 | 0.802 | 0.69 |
| **orf10c** | 0.944 | 0.695 | 0.725 | 0.805 | 0.734 | 0.866 | 0.824 | 0.714 | 0.807 | 0 | 0.734 | 0.851 | 0.896 | 0.635 | 0.86 | 0.757 | 0.843 |
| **lovF** | 0.899 | 0.807 | 0.603 | 0.771 | 0.708 | 0.86 | 0.824 | 0.755 | 0.825 | 0.734 | 0 | 0.823 | 0.9 | 0.837 | 0.91 | 0.878 | 0.808 |
| **orf13** | 0.884 | 0.806 | 0.839 | 0.904 | 0.916 | 0.862 | 0.949 | 0.669 | 0.784 | 0.851 | 0.823 | 0 | 0.958 | 0.703 | 0.762 | 0.605 | 0.812 |
| **orf14** | 0.899 | 0.951 | 0.924 | 0.827 | 0.901 | 0.945 | 1 | 0.95 | 0.786 | 0.896 | 0.9 | 0.958 | 0 | 0.947 | 0.952 | 0.953 | 0.855 |
| **orf15** | 0.883 | 0.629 | 0.859 | 0.916 | 0.889 | 0.882 | 0.931 | 0.674 | 0.712 | 0.635 | 0.837 | 0.703 | 0.947 | 0 | 0.739 | 0.74 | 0.789 |
| **orf16** | 0.918 | 0.83 | 0.96 | 0.81 | 0.915 | 0.848 | 0.97 | 0.495 | 0.808 | 0.86 | 0.91 | 0.762 | 0.952 | 0.739 | 0 | 0.607 | 0.787 |
| **orf17** | 0.912 | 0.705 | 0.857 | 0.829 | 0.922 | 0.852 | 0.944 | 0.557 | 0.802 | 0.757 | 0.878 | 0.605 | 0.953 | 0.74 | 0.607 | 0 | 0.769 |
| **orf18** | 0.804 | 0.773 | 0.791 | 0.888 | 0.727 | 1 | 0.871 | 0.722 | 0.69 | 0.843 | 0.808 | 0.812 | 0.855 | 0.789 | 0.787 | 0.769 | 0 |

**Table S6.** Evolutionary distance matrix of the Lovastatin BGC of *Aspergillus terreus* (lov)

| | orf1 | orf2 | lovA | lovB | lovG | lovC | lovD | orf80c | lovE | orf10c | lovF | orf13 | orf14 | orf15 | orf16 | orf17 | orf18 |
|---|---|---|---|---|---|---|---|---|---|---|---|---|---|---|---|---|---|
| orf1 | 0 | 0.434 | 0.16 | 0.301 | 0.517 | 0.289 | 0.389 | 0.495 | 0.484 | 0.858 | 0.16 | 0.211 | 0.484 | 0.821 | 0.494 | 0.525 | 0 |
| orf2 | 0.434 | 0 | 0.247 | 0 | 0.329 | 0.07 | 0.301 | 0.205 | 0.326 | 0 | 0 | 0.118 | 0.394 | 0 | 0.152 | 0 | 0 |
| lovA | 0.16 | 0.247 | 0 | 0.161 | 0.202 | 0.774 | 0 | 0.445 | 0 | 0 | 0 | 0 | 0.817 | 0 | 0.72 | 0 | 0.32 |
| lovB | 0.301 | 0 | 0.161 | 0 | 0 | 0.394 | 0 | 0.12 | 0.088 | 0.349 | 0.16 | 0.088 | 0.133 | 0.449 | 0 | 0 | 0.284 |
| lovG | 0.517 | 0.329 | 0.202 | 0 | 0 | 0.34 | 0 | 0.478 | 0.629 | 0.262 | 0.202 | 0.255 | 0.477 | 0.439 | 0.452 | 0.451 | 0.294 |
| lovC | 0.289 | 0.07 | 0.774 | 0.394 | 0.34 | 0 | 0.5 | 0 | 0.333 | 0.293 | 0.123 | 0 | 0.75 | 0.184 | 0 | 0 | 1 |
| lovD | 0.389 | 0.301 | 0 | 0 | 0 | 0.5 | 0 | 0.679 | 1 | 0.378 | 0 | 0 | 1 | 0 | 0.706 | 0 | 0.25 |
| orf80c | 0.495 | 0.205 | 0.445 | 0.12 | 0.478 | 0 | 0.679 | 0 | 0 | 0.393 | 0.374 | 0 | 0.805 | 0.173 | 0 | 0 | 0 |
| lovE | 0.484 | 0.326 | 0 | 0.088 | 0.629 | 0.333 | 1 | 0 | 0 | 0.485 | 0 | 0 | 0 | 0.292 | 0 | 0 | 0.208 |
| orf10c | 0.858 | 0 | 0 | 0.333 | 0.262 | 0.3 | 0.378 | 0.393 | 0.485 | 0 | 0 | 0.393 | 0.485 | 0 | 0.253 | 0.301 | 0.147 |
| lovF | 0.16 | 0 | 0 | 0.16 | 0.202 | 0.123 | 0 | 0.374 | 0 | 0 | 0 | 0 | 0 | 0 | 0.175 | 0 | 0 |
| orf13 | 0.211 | 0.118 | 0 | 0.088 | 0.282 | 0 | 0 | 0 | 0 | 0.393 | 0 | 0 | 0 | 0 | 0 | 0 | 0 |
| orf14 | 0.484 | 0.394 | 0.817 | 0.133 | 0.477 | 0.75 | 1 | 0.805 | 0 | 0.485 | 0 | 0 | 0 | 0.615 | 0.823 | 0 | 0 |
| orf15 | 0.821 | 0 | 0 | 0.449 | 0.439 | 0.184 | 0 | 0.173 | 0.292 | 0 | 0 | 0 | 0.615 | 0 | 0 | 0.171 | 0.192 |
| orf16 | 0.494 | 0.152 | 0.72 | 0 | 0.452 | 0 | 0.706 | 0 | 0 | 0.253 | 0.175 | 0 | 0.823 | 0 | 0 | 0 | 0 |
| orf17 | 0.525 | 0 | 0 | 0 | 0.451 | 0 | 0 | 0 | 0 | 0.245 | 0 | 0 | 0 | 0.171 | 0 | 0 | 0 |
| orf18 | 0 | 0 | 0.32 | 0.284 | 0.294 | 1 | 0.25 | 0 | 0.208 | 0.147 | 0 | 0 | 0 | 0.192 | 0 | 0 | 0 |

**Table S7.** Combined distance matrix of the Lovastatin BGC of *Aspergillus terreus* (lov)

| | orf1 | orf2 | lovA | lovB | lovG | lovC | lovD | orf80c | lovE | orf10c | lovF | orf13 | orf14 | orf15 | orf16 | orf17 | orf18 |
|---|---|---|---|---|---|---|---|---|---|---|---|---|---|---|---|---|---|
| orf1 | 0 | 1,287 | 1,075 | 1,158 | 1,417 | 1,18 | 1,321 | 1,404 | 1,29 | 1,802 | 1,059 | 1,095 | 1,383 | 1,704 | 1,412 | 1,437 | 0,804 |
| orf2 | 1,287 | 0 | 1,12 | 0,824 | 1,193 | 0,915 | 1,205 | 0,88 | 1,137 | 0,695 | 0,807 | 0,924 | 1,345 | 0,629 | 0,982 | 0,705 | 0,773 |
| lovA | 1,075 | 1,12 | 0 | 0,99 | 0,913 | 1,692 | 0,738 | 1,318 | 0,844 | 0,725 | 0,603 | 0,839 | 1,741 | 0,859 | 1,68 | 0,857 | 1,111 |
| lovB | 1,158 | 0,824 | 0,99 | 0 | 0,57 | 1,054 | 0,746 | 0,99 | 0,851 | 1,154 | 0,931 | 0,992 | 0,96 | 1,365 | 0,81 | 0,829 | 1,172 |
| lovG | 1,417 | 1,193 | 0,913 | 0,57 | 0 | 1,153 | 0,845 | 1,385 | 1,346 | 0,996 | 0,91 | 1,171 | 1,378 | 1,328 | 1,367 | 1,373 | 1,021 |
| lovC | 1,18 | 0,915 | 1,692 | 1,054 | 1,153 | 0 | 1,363 | 0,834 | 1,141 | 1,159 | 0,983 | 0,862 | 1,695 | 1,066 | 0,848 | 0,852 | 2 |
| lovD | 1,321 | 1,205 | 0,738 | 0,746 | 0,845 | 1,363 | 0 | 1,647 | 2 | 1,202 | 0,824 | 0,949 | 2 | 0,931 | 1,676 | 0,944 | 1,121 |
| orf80c | 1,404 | 0,88 | 1,318 | 0,99 | 1,385 | 0,834 | 1,647 | 0 | 0,802 | 1,107 | 1,129 | 0,669 | 1,755 | 0,847 | 0,495 | 0,557 | 0,722 |
| lovE | 1,29 | 1,137 | 0,844 | 0,851 | 1,346 | 1,141 | 2 | 0,802 | 0 | 1,292 | 0,825 | 0,784 | 0,786 | 1,004 | 0,808 | 0,802 | 0,898 |
| orf10c | 1,802 | 0,695 | 0,725 | 1,138 | 0,996 | 1,166 | 1,202 | 1,107 | 1,292 | 0 | 0,734 | 1,244 | 1,381 | 0,635 | 1,113 | 1,058 | 0,99 |
| lovF | 1,059 | 0,807 | 0,603 | 0,931 | 0,91 | 0,983 | 0,824 | 1,129 | 0,825 | 0,734 | 0 | 0,823 | 0,9 | 0,837 | 1,085 | 0,878 | 0,808 |
| orf13 | 1,095 | 0,924 | 0,839 | 0,992 | 1,198 | 0,862 | 0,949 | 0,669 | 0,784 | 1,244 | 0,823 | 0 | 0,958 | 0,703 | 0,762 | 0,605 | 0,812 |
| orf14 | 1,383 | 1,345 | 1,741 | 0,96 | 1,378 | 1,695 | 2 | 1,755 | 0,786 | 1,381 | 0,9 | 0,958 | 0 | 1,562 | 1,775 | 0,953 | 0,855 |
| orf15 | 1,704 | 0,629 | 0,859 | 1,365 | 1,328 | 1,066 | 0,931 | 0,847 | 1,004 | 0,635 | 0,837 | 0,703 | 1,562 | 0 | 0,739 | 0,911 | 0,981 |
| orf16 | 1,412 | 0,982 | 1,68 | 0,81 | 1,367 | 0,848 | 1,676 | 0,495 | 0,808 | 1,113 | 1,085 | 0,762 | 1,775 | 0,739 | 0 | 0,607 | 0,787 |
| orf17 | 1,437 | 0,705 | 0,857 | 0,829 | 1,373 | 0,852 | 0,944 | 0,557 | 0,802 | 1,002 | 0,878 | 0,605 | 0,953 | 0,911 | 0,607 | 0 | 0,769 |
| orf18 | 0,804 | 0,773 | 1,111 | 1,172 | 1,021 | 2 | 1,121 | 0,722 | 0,898 | 0,99 | 0,808 | 0,812 | 0,855 | 0,981 | 0,787 | 0,769 | 0 |

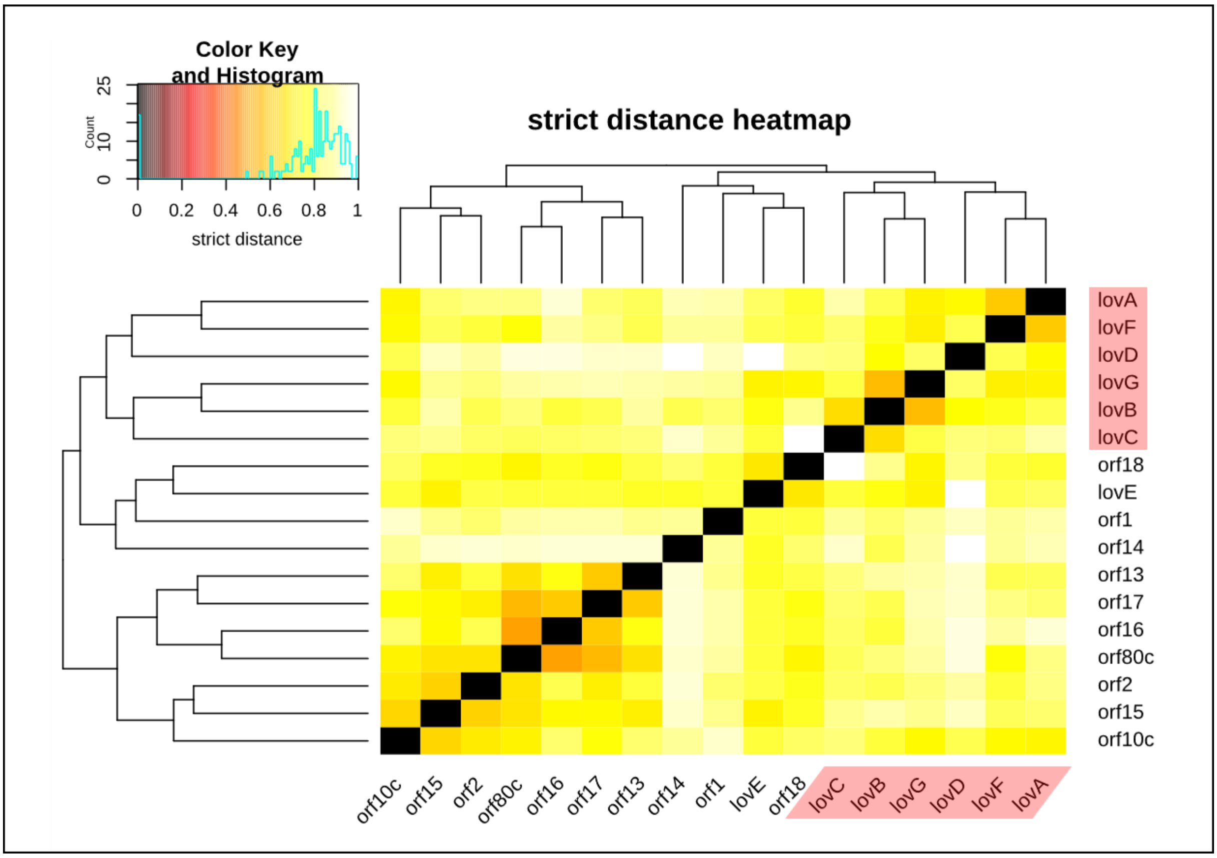


**Figure S1.** Heatmap of the strict distance matrix of the Lovastatin BGC of *Aspergillus terreus* (lov). This is part of the standard output of the FunOrder analysis. The genes necessary for the biosynthesis of lovastatin are highlighted in red.

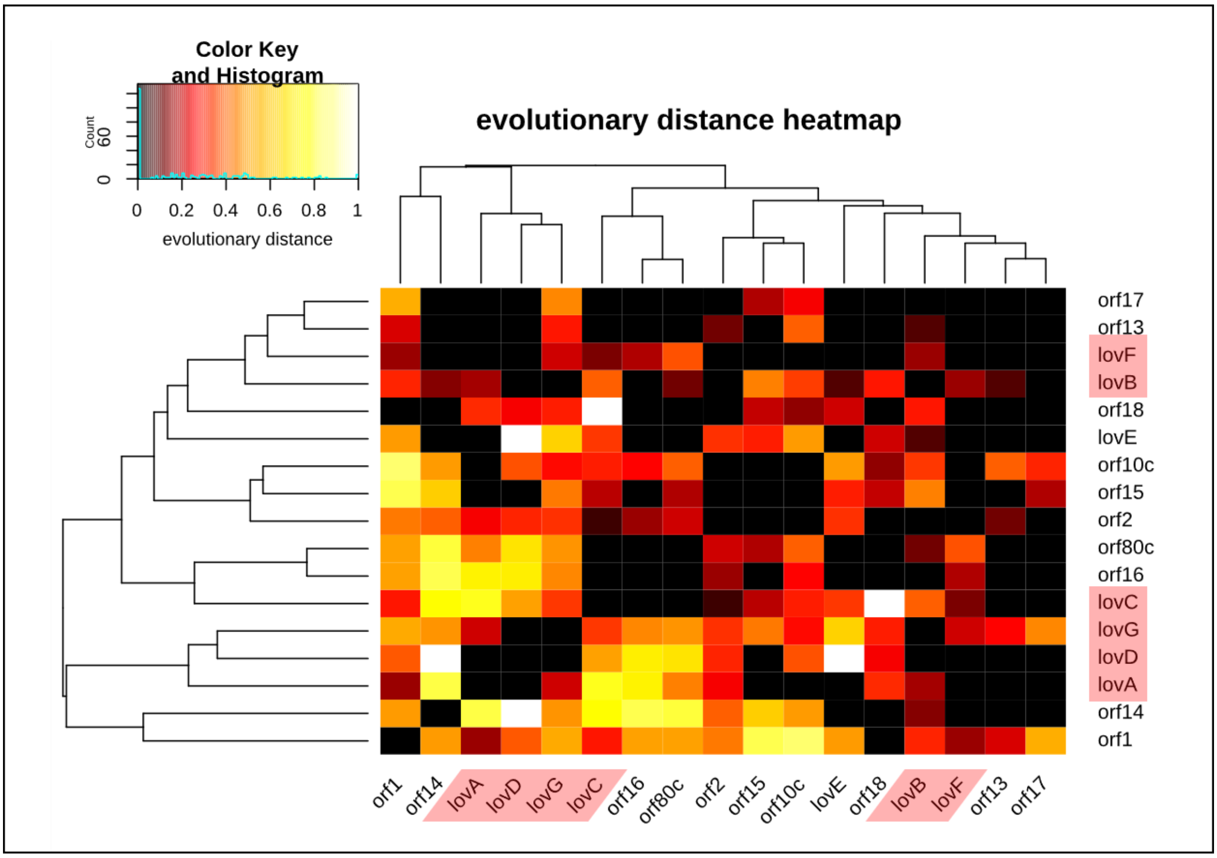


**Figure S2.** Heatmap of the evolutionary distance matrix of the Lovastatin BGC of *Aspergillus terreus* (lov). This is part of the standard output of the FunOrder analysis. The genes necessary for the biosynthesis of lovastatin are highlighted in red.

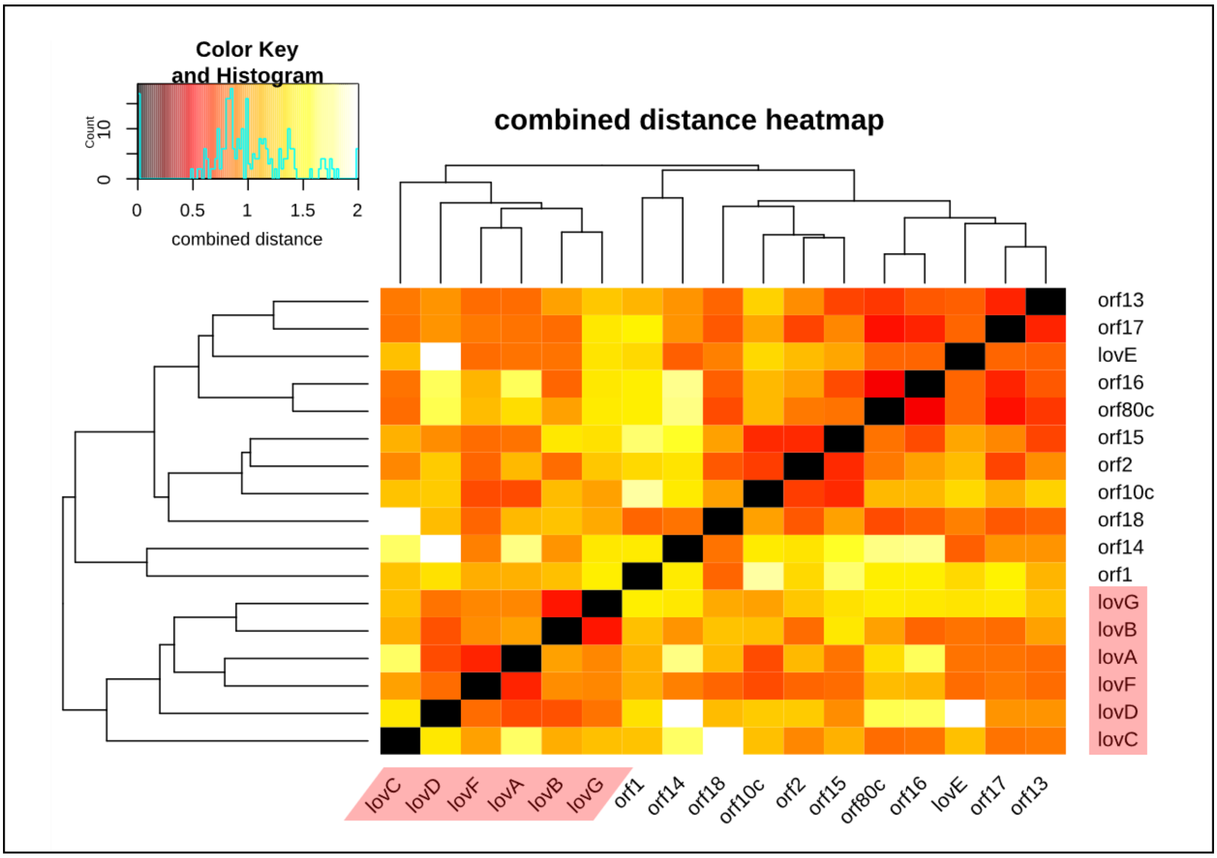


**Figure S3.** Heatmap of the combined distance matrix of the Lovastatin BGC of *Aspergillus terreus* (lov). This is part of the standard output of the FunOrder analysis. The genes necessary for the biosynthesis of lovastatin are highlighted in red.

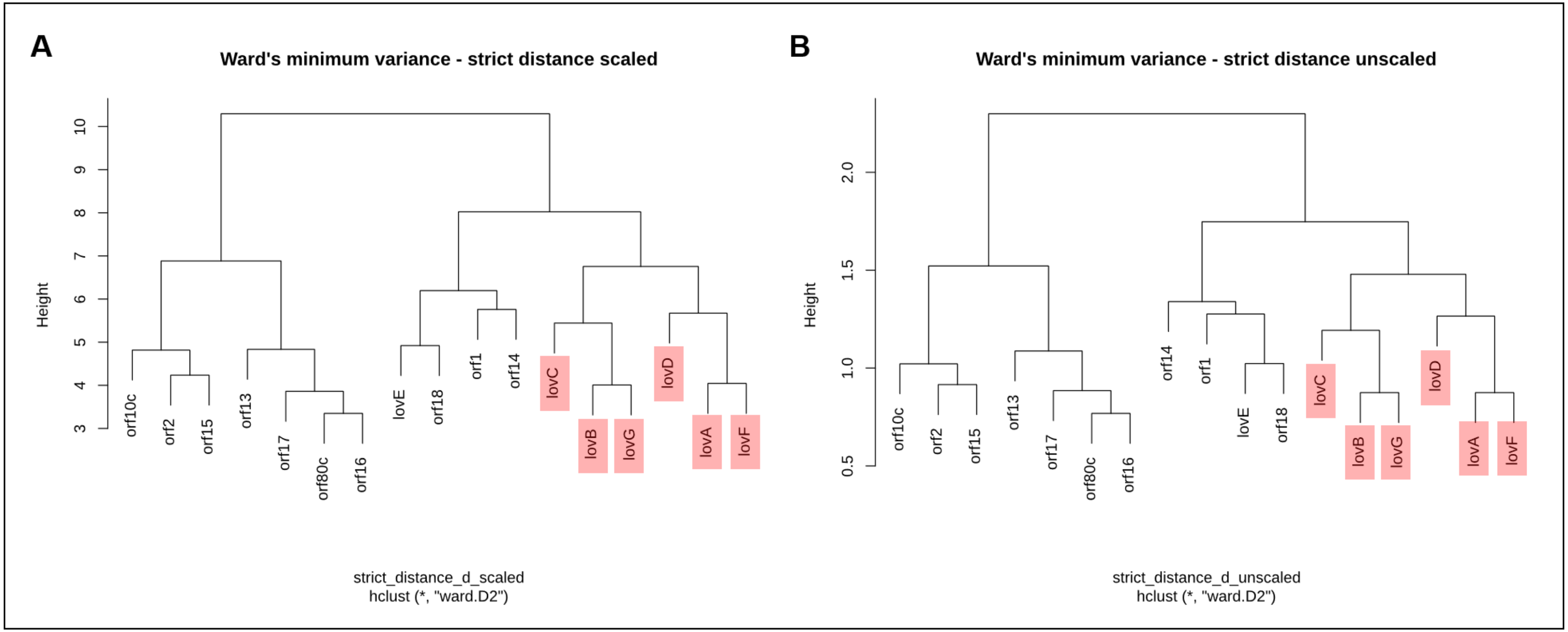


**Figure S4.** Dendrograms based on the scaled (A) and unscaled (B) strict distance matrix of the Lovastatin BGC of *Aspergillus terreus* (lov). The clustering was performed based on the Euclidean distance within the matrix using Ward's minimum variance method aiming at finding compact spherical clusters, with the implemented squaring of the dissimilarities before cluster updating. This is part of the standard output of the FunOrder analysis. The genes necessary for the biosynthesis of lovastatin are highlighted in red.

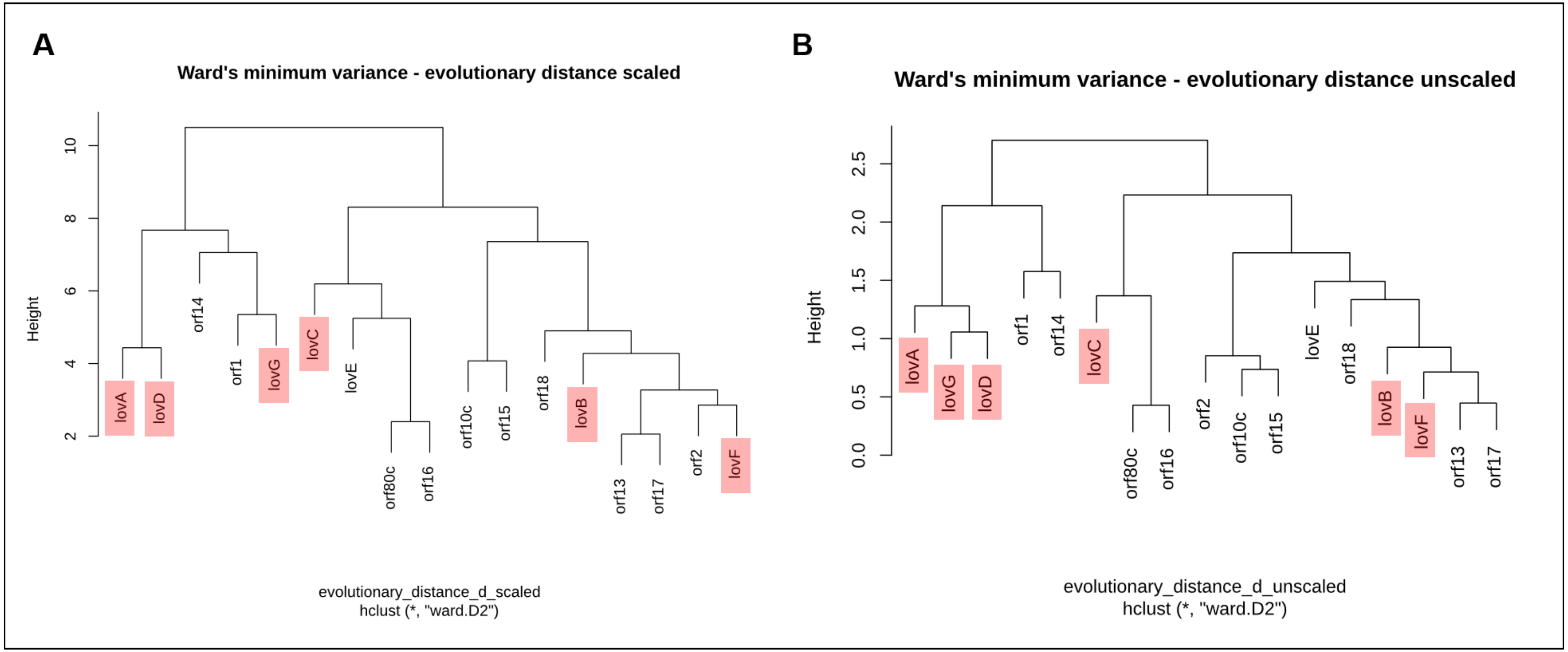


**Figure S5.** Dendrograms based on the scaled (A) and unscaled (B) evolutionary distance matrix of the Lovastatin BGC of *Aspergillus terreus* (lov). The clustering was performed based on the Euclidean distance within the matrix using Ward's minimum variance method aiming at finding compact spherical clusters, with the implemented squaring of the dissimilarities before cluster updating. This is part of the standard output of the FunOrder analysis. The genes necessary for the biosynthesis of lovastatin are highlighted in red.

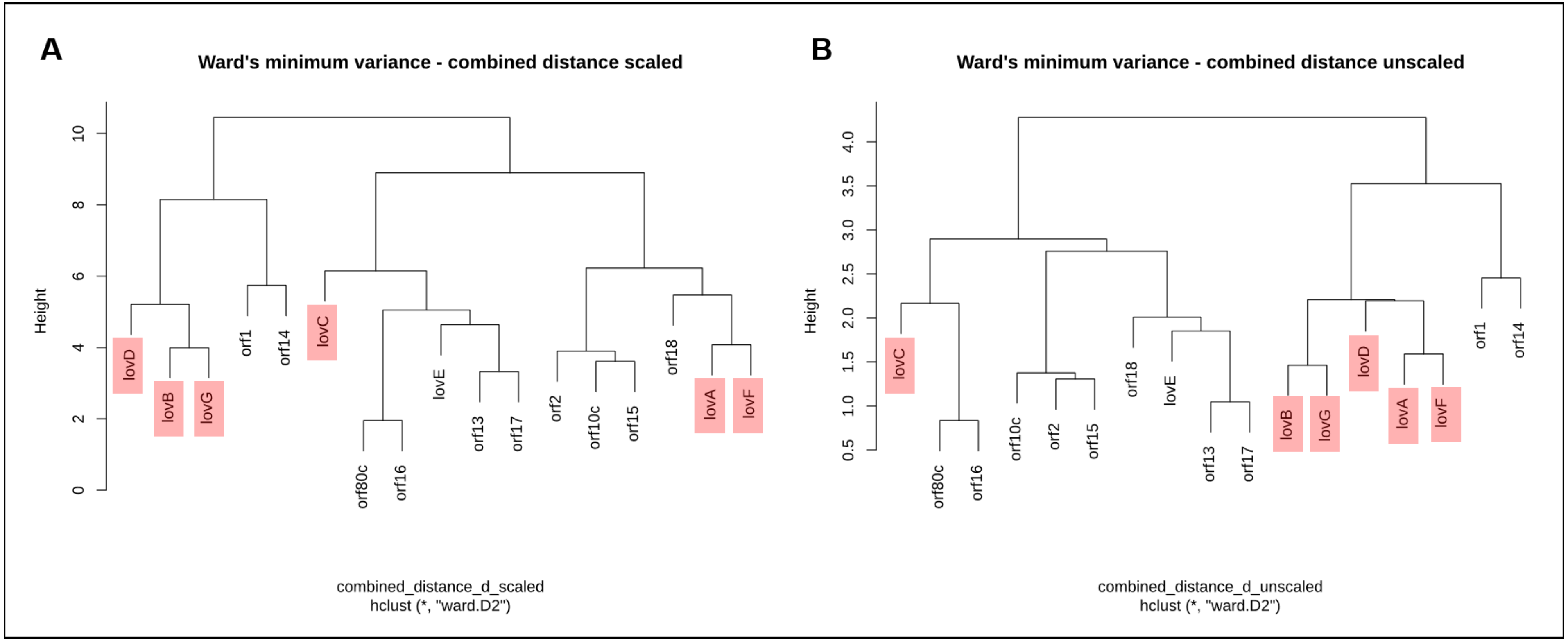


**Figure S6.** Dendrograms based on the scaled (A) and unscaled (B) combined distance matrix of the Lovastatin BGC of *Aspergillus terreus* (lov). The clustering was performed based on the Euclidean distance within the matrix using Ward's minimum variance method aiming at finding compact spherical clusters, with the implemented squaring of the dissimilarities before cluster updating. This is part of the standard output of the FunOrder analysis. The genes necessary for the biosynthesis of lovastatin are highlighted in red.

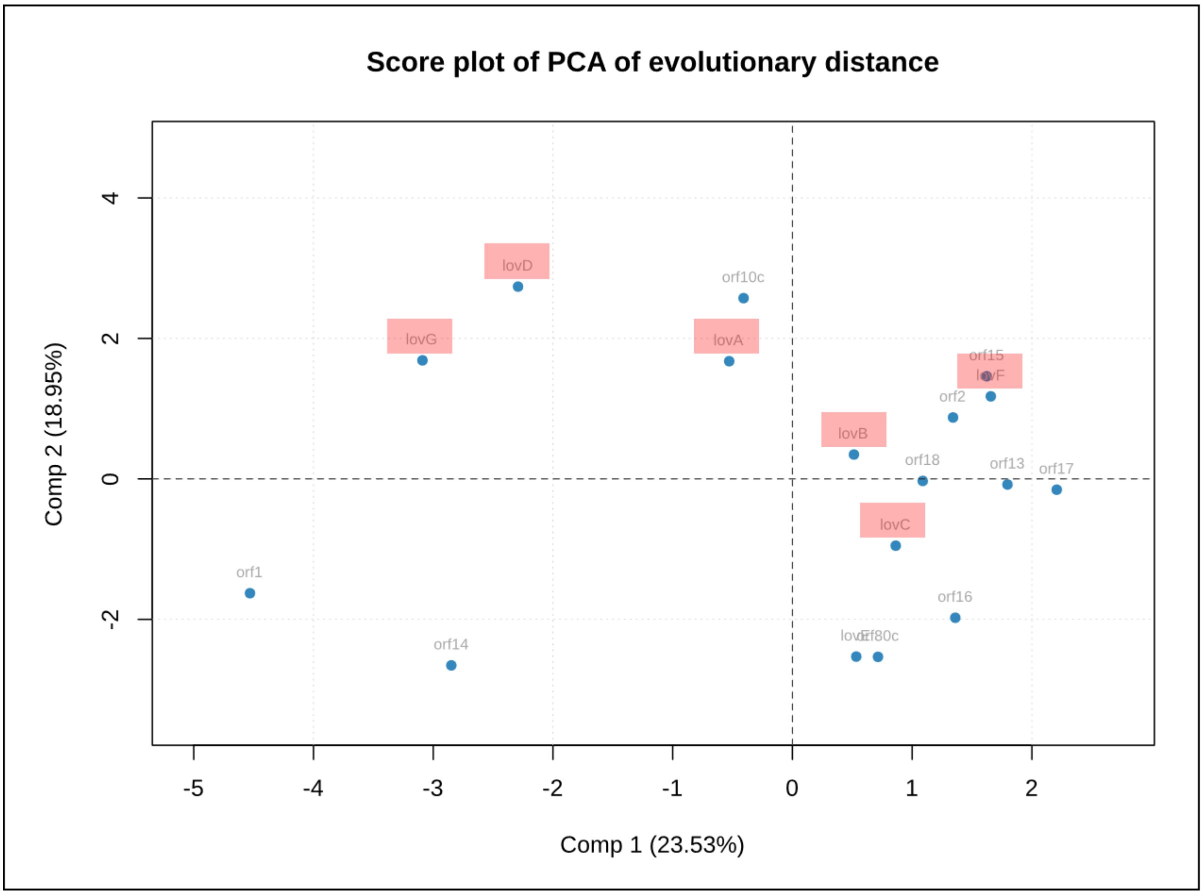


**Figure S7.** Score plot of the first two principal components from a PCA performed on the evolutionary distance matrix analysis of the Lovastatin BGC of *Aspergillus terreus* (lov). This is part of the standard output of the FunOrder analysis. The genes necessary for the biosynthesis of lovastatin are highlighted in red.
